# Development and application of GlycanDIA workflow for glycomic analysis

**DOI:** 10.1101/2024.03.12.584702

**Authors:** Yixuan Xie, Xingyu Liu, Chenfeng Zhao, Siyu Chen, Shunyang Wang, Zongtao Lin, Faith M. Robison, Benson M. George, Ryan A. Flynn, Carlito B. Lebrilla, Benjamin A. Garcia

## Abstract

Glycans modify protein, lipid, and even RNA molecules to form the regulatory outer coat on cells called the glycocalyx. The changes in glycosylation have been linked to the initiation and progression of many diseases. Thus, while the significance of glycosylation is well established, a lack of accessible methods to characterize glycans has hindered the ability to understand their biological functions. Mass spectrometry (MS)-based methods have generally been at the core of most glycan profiling efforts; however, modern data-independent acquisition (DIA), which could increase sensitivity and simplify workflows, has not been benchmarked for analyzing glycans. Herein, we developed a DIA-based glycomic workflow, termed GlycanDIA, to identify and quantify glycans with high sensitivity and accuracy. The GlycanDIA workflow combined higher energy collisional dissociation (HCD)-MS/MS and staggered windows for glycomic analysis, which facilitates the sensitivity in identification and the accuracy in quantification compared to conventional data-dependent acquisition (DDA)-based glycomics. To facilitate its use, we also developed a generic search engine, GlycanDIA Finder, incorporating an iterative decoy searching for confident glycan identification and quantification from DIA data. The results showed that GlycanDIA can distinguish glycan composition and isomers from *N*-glycans, *O*-glycans, and human milk oligosaccharides (HMOs), while it also reveals information on low-abundant modified glycans. With the improved sensitivity, we performed experiments to profile *N*-glycans from RNA samples, which have been underrepresented due to their low abundance. Using this integrative workflow to unravel the *N*-glycan profile in cellular and tissue glycoRNA samples, we found that RNA-glycans have specific forms as compared to protein-glycans and are also tissue-specific differences, suggesting distinct functions in biological processes. Overall, GlycanDIA can provide comprehensive information for glycan identification and quantification, enabling researchers to obtain in-depth and refined details on the biological roles of glycosylation.

**Graphical Abstract:** 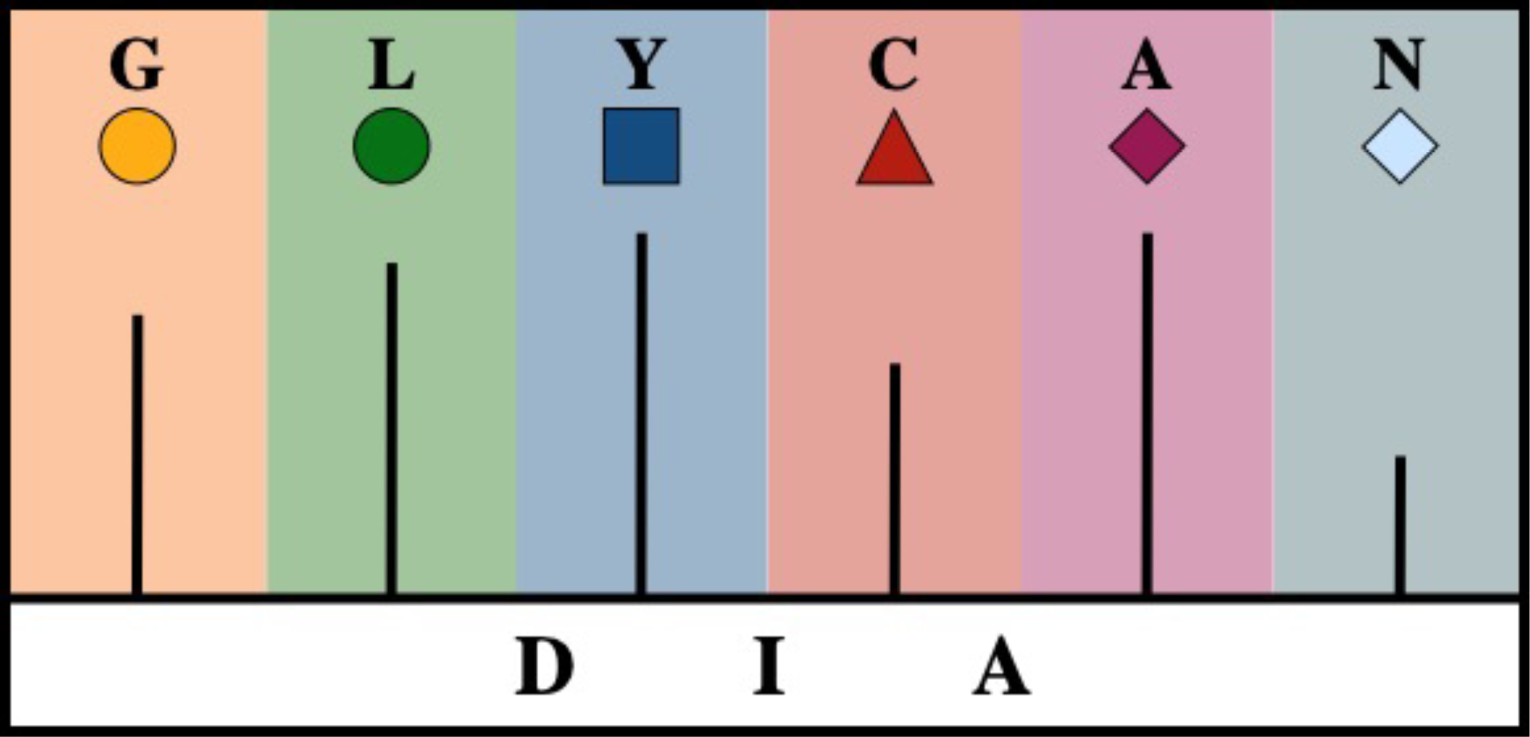

## Introduction

Glycosylation is a major post-translational modification (PTM) of proteins and lipids, and the glycosylated RNAs were also recently found on the cell membrane.^1–3^ Glycans are fundamental for cellular functional processes, including cell adhesion, cell signaling, immunological response, and cancer metastasis.^4^ The extension of oligosaccharide chains involves a complicated competition between various glycosyltransferases, leading to the observed heterogeneity of glycan structures. Glycans are historically analyzed using nuclear magnetic resonance (NMR) or lectin-array.^5, 6^ However, the high sample demand or the lack of sequence information for each glycan is not ideal for comprehensive analysis. Therefore, developing new techniques to characterize the glycan chains is essential to understanding the roles of glycosylation in cell physiology.

Mass spectrometry (MS)-based methods emerged as a powerful tool and have significantly advanced glycomic analysis, providing information about glycan composition, structure, and abundance.^7^ The biosynthetic pathway of glycans often yields glycan isomers, and tandem MS is required for their differentiation and determination. Data-dependent acquisition (DDA) has been the predominant approach for resolving glycan information, wherein only the most abundant precursor ions (top N) are selected for tandem MS/MS (MS2) analysis (**Figure 1A**).^8, 9^ This feature from DDA-based methods often leads to underrepresented and inconsistent detection of low-abundance molecules. This is particularly important in material-limited contexts with clinical samples or places where glycan levels are low like RNA. For example, to investigate the glycan alterations during embryonic development, millions of sorted embryonic cells are collected, while limited glycan information is acquired using the DDA-based method.^10^ To improve the detection, native N-glycans often need labeling or derivatization to decrease sample complexity and/or enhance ionization efficiency in DDA, while it requires extra processing steps and generates issues in isomer separation.^11^ To tackle these challenges, targeted analysis, such as multiple reaction monitoring (MRM), has been employed for glycomics, which relies on the product ions from different glycans and provides reliable quantitative results due to higher specificity and less variation of the fragment ions.^12^ However, only specific glycoforms of interest can be monitored, which is not optimal for glycan discovery in novel samples such as glycosylated

**Figure 1.**
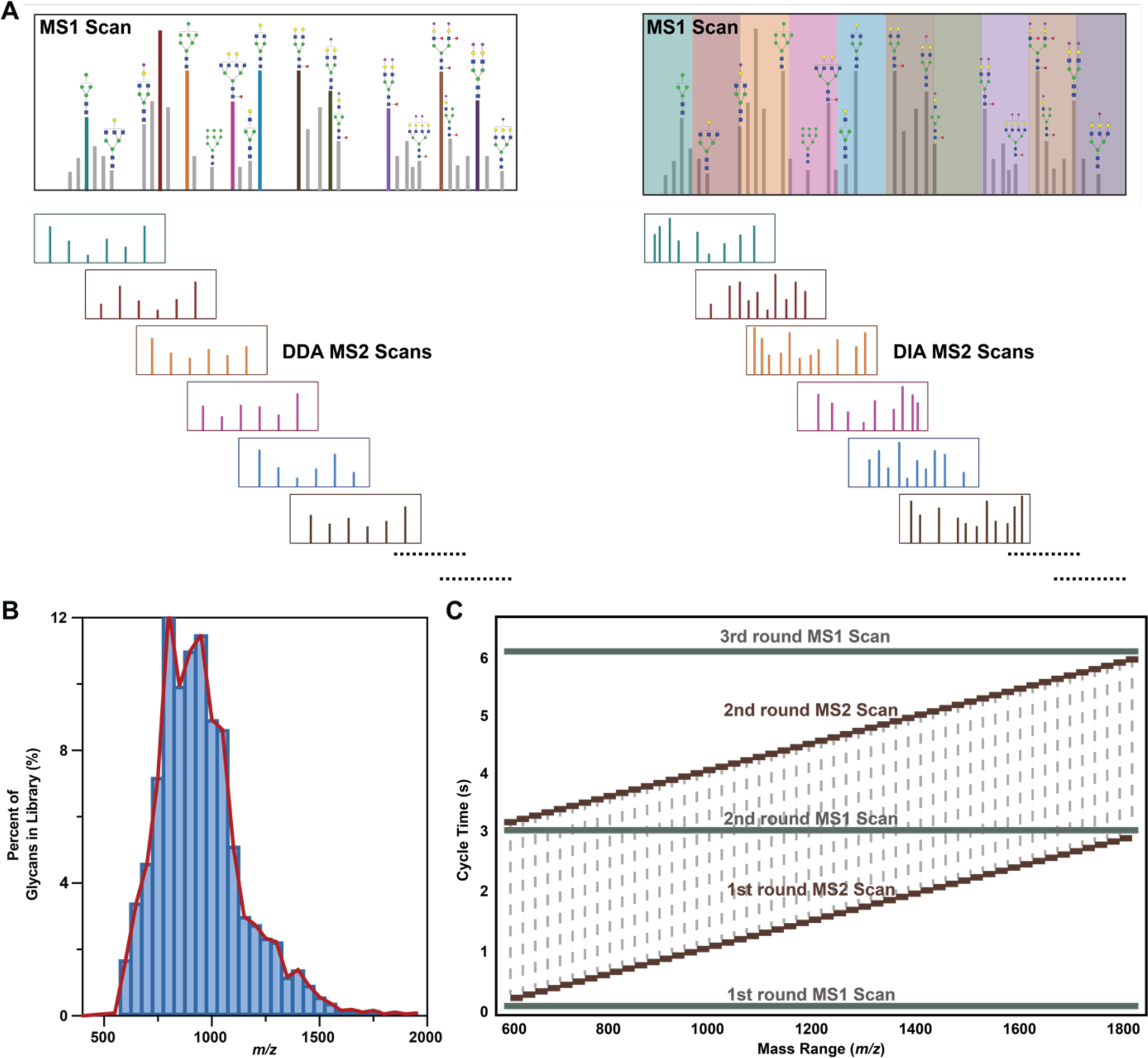
MS analysis in GlycanDIA workflow. **A.** In the conventional data-dependent acquisition (DDA)-based method (left), a few most abundant precursor ions are respectively selected within a small mass window (typically 0.7-2 *m/z*) and are further fragmented. DDA ideally only generates the tandem MS/MS (MS2) information for specific ions. During the data-independent acquisition (DIA) (right), pre-defined wide isolation windows are applied without regard to the abundance of precursor ions, and all precursor ions within the isolation window are fragmented in MS2. DIA often yields mixed MS2 information from different ions. **B.** To cover all the *N*-glycans using GlycanDIA, the released *N*-glycans from different cells were mapped, and all their masses were within the range of 500-2000 *m/z*. Importantly, the precursor masses from 600 to 1800 *m/z* cover all glycan species. **C.** The illustration of staggered windowing schemes in GlycanDIA. The first cycle consisted of one MS1 scan and different DIA scans across the mass range of selection. The second cycle consisted of another MS1 scan and different DIA scans that staggered the first-round window. Dashed lines indicate when an isolation window range is repeated.

RNA. Data-independent acquisition (DIA)-based methods have been introduced, which do not rely on precursor ion selection;^13^ instead, all precursors within a predefined mass window are fragmented simultaneously. Because DIA can generate an unbiased and comprehensive dataset, it has been more recently widely applied to the characterization of peptides, lipids, metabolites, and nucleosides.^14–16^ However, due to the complexity of glycan structures, lacking specific instrument methods, and the bioinformatic tools were barely designed for searching glycomics data, to our knowledge, the application of DIA for glycomic analysis has not been established.

Herein, we analyzed glycans in the positive mode to establish a DIA-based glycomic workflow, termed GlycanDIA, utilizing higher energy collisional dissociation (HCD)-MS/MS and staggered DIA windows for glycomic analysis. We analyzed the data using a combination of both MS1-centric and MS2-centric strategies while further developing a generic search engine, GlycanDIA Finder, to perform automated data analysis with iterative decoy searching. This workflow can enhance the identification of glycoforms and improve the accuracy of glycan quantification compared to DDA. We have demonstrated that GlycanDIA workflow enables confident identification of *N*-glycans, *O*-glycans, and human milk oligosaccharides (HMOs). We have also discriminated different glycan isomers (including composition and linkage isomers) and elucidated the profile of low-abundant modified species using GlycanDIA. Such low-level species include *N*-glycans on RNAs, which were recently discovered but are not fully profiled due to their extremely low abundance. To demonstrate the advantage of the GlycanDIA workflow, we applied this new approach to measuring these *N*-glycans on RNA extracted from cultured human cell lines and mouse tissues. This application yielded comprehensive and quantitative information about the *N*-glycan landscape of RNA extracts from different samples. This GlycanDIA workflow can be broadly applied for monitoring glycosylation status in different biological conditions, providing researchers with a comprehensive view of glycobiology.

## Results

### Establishment of GlycanDIA workflow

Electrospray ionization is amenable for neutral and anionic native glycans in the positive mode, and to find the optimal fragmentation energy for the analysis, we first optimized the normalized collision energy (NCE) for HCD fragmentation. Examples shown in **Figure S1-3**, the sequencing ions of the represented high mannose, fucosylated, and sialylated glycans first increased with rising NCE due to more efficient fragmentation of the precursor ions, while the larger fragment ions decreased when the collision energy was set to be greater than 25% due to over-fragmentation. As a result, 20% NCE was selected as the fragmentation energy for generating the best sequence information for the majority of *N*-glycans. Notably, although cross-ring fragments can be generated from glycans, the majority of fragment ions (>99.5%) were generated from glycosidic bond cleavages (**Figure S4**). This result is in agreement with previous observations.^17^

DIA-based methods fragment all precursor ions from an isolation window, generating highly multiplexed fragment ion spectra. It was therefore critical to optimize the mass window scheme for the glycan analysis. We mapped the previously profiled *N*-glycans from eight cell lines that employed DDA-MS to acquire the general *N*-glycan distributions.^18, 19^ As shown in **Figure 1B**, we found that the *N*-glycan precursors allocate to a wide mass-to-charge range (500-2000 *m/z*), while the precursor range of 600-1800 *m/z* covers all different major ion species. After selecting the precursor window, we evaluated different strategies to set up the mass window scheme, including fixed DIA, staggered DIA, multiplexed (MSX) DIA, and variable DIA. We balanced the size of the isolation window and the cycle of scans (the total counts of isolation windows) in different DIA strategies. As shown in **Figure S5-8**, staggered and variable DIA showed better results in generating fewer interfering ions. It was also noted that the loop counts (window numbers) for GlycanDIA are larger than that of the common DIA for proteomics. This is due to the full width at half maximum (FWHM) of most glycans on the PGC column being more than 0.3 minutes (**Figure S9A**). As a result, most glycan compounds had sufficient data points (∼10) for constructing Gaussian peaks (**Figure S9B**) using the 24 *m/z* staggered method, which yielded higher accuracy for quantification. The representative scan scheme of staggered windowing and detailed parameters are shown in **Figure 1C** and **Table S1**, respectively. Meanwhile, although the variable windowing showed decent results in quantification, the method was cumbersome to set up and adjust using the manufacturer’s software; therefore, we used 24 *m/z* staggered with 50 windows for further analysis.

In DDA-MS, only one precursor is theoretically picked and fragmented per MS/MS scan, whereas DIA contains complicated MS2 information due to the collection of fragmentation from all coeluting compounds within a specified mass window. To decipher the glycan information from the DIA data, we used the MS1-centric method in the GlycanDIA workflow (**Figure 2A**). Taking the *N*-glycan Hex(4)HexNAc(4)Fuc(0)Neu5Ac(1) (4401 for short) as an example, the possible precursor ion masses were calculated and specifically extracted from the MS1 level. After locating the peak, the product ion yielded from this glycan (*e.g.*, 1258.45 *m/z*) was extracted from two MS2 spectra obtained from the window containing the precursor ion (876-900 *m/z* and 864-888 *m/z*) to confirm the fragmentation and one spectrum from a nearby window (888-912 *m/z*) as a negative control. As a result, only the 876-900 *m/z* and 864-888 *m/z* windows showed extracted product ions coeluting with the precursor ion, and multiple product ions were extracted simultaneously to validate the sequence of the glycan. In addition, we also employed a complementary MS2-centric strategy to find the glycans via glycan signature product ions from a random MS2 spectrum. For example, 292.10 *m/z* standing for Neu5Ac was first extracted from the 864-888 *m/z* window. After examining the same ion from the two adjacent MS2 spectra, 876-900 *m/z* and 852-876 *m/z*, we found that only the MS2 from 876-900 *m/z* contained the 292.10 *m/z* fragment ion and coeluted with the 864-888 *m/z* window. This result means the precursor of this product ion fell into the overlap of the two windows, 876-888 *m/z*. After extracting the MS1 spectrum from 876 to 888 *m/z*, the doubly charged monoisotopic ion at 885.8257 *m/z* was noted, and the glycan composition, 4401, can be determined and validated by revisiting the MS2 information. Collectively, the two peaks standing for Hex(4)HexNAc(4)Fuc(0)Neu5Ac(1) anomers were identified (**Figure 2B**), and the two approaches were used at the same time to cross-validate and fully resolve the glycan information yielded from DIA data.

**Figure 2.**
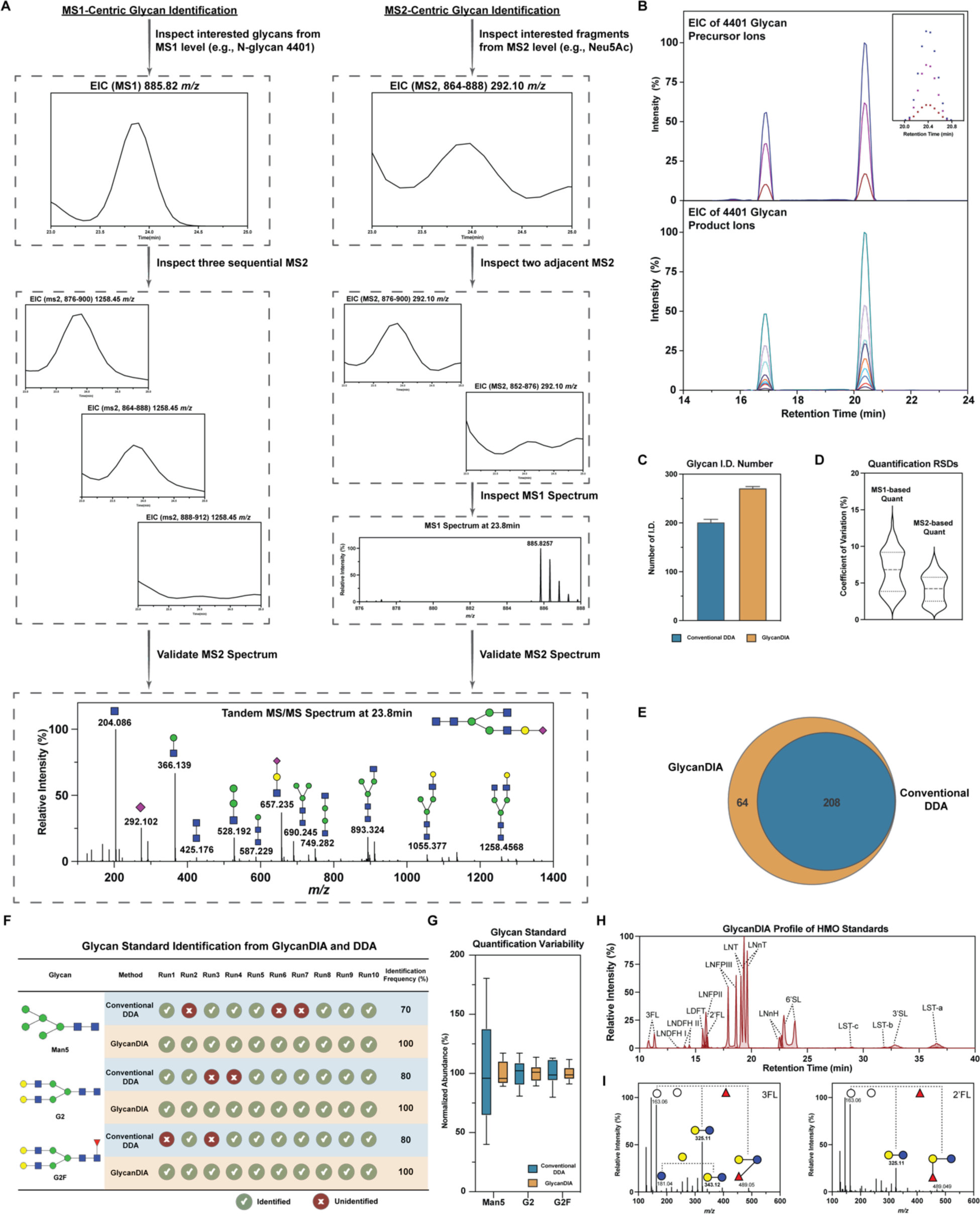
The results from GlycanDIA workflow. **A.** To analyze GlycanDIA data, the MS1-centric method needs firstly to calculate the precursor ion mass and monitor it from the MS1 level (885.82 *m/z* in this example). Subsequently, the product ion yielded from this glycan (e.g., 1258.45 *m/z*) was also monitored from the MS2 spectra that isolated the precursor ion. A series of product ions were extracted simultaneously to validate the glycan sequence. The MS2-centric strategy in the GlycanDIA data analysis strategy relied on the glycan signature ions from a random MS2 spectrum. In this example, the fragment of 292.10 *m/z* from sialylated glycans was monitored from three adjacent MS2 spectra. The peak only aligned from the two corresponding windows, indicating the product was from the precursor masses within the mass range that overlapped with the two specified windows. The glycan composition can be further confirmed by validating MS1 and MS2 information. **B.** The peak alignment between glycan precursor ions at the MS1 level (left) and glycan product ions at the MS2 level (right) suggests the unambiguous identification of glycans from GlycanDIA data. The small box figure shows sufficient data points across the peak for quantification. **C.** More than 270 different *N*-glycans were identified using GlycanDIA, while the conventional DDA-based method only identified 200 glycans. **D.** The glycan quantification in MS1 and MS2 levels was evaluated, and the quantitative results using MS2 information can generate fewer variations. **E.** GlycanDIA covered all the glycans identified from DDA runs. These suggest the improved ability of GlycanDIA to identify glycans. **F.** Three heavy isotope-labeled *N*-glycan standards were mixed with the *N*-glycan pool released from RNase B and human sera. The GlycanDIA provides more consistent glycan identification compared to the DDA-based method. **G.** The GlycanDIA also generates less variations in quantitive results. **H.** Extracted compound chromatogram (ECC) shows the elution profile of 15 HMO standards. The full name of HMO standards can be found in **Table S2**. **I.** The example GlycanDIA results of two HMO isomers, 3FL and 2’FL. The results show GlycanDIA disclosing fragmentation information that enables the distinguishment of glycan isomers.

### Development of GlycanDIA Finder as a computational software

To meet the need for computational tools that streamline data interpretation and extend the application of the workflow, we developed a generic search engine, GlycanDIA Finder, upon Python and MatchMS library.^20^ This in-house developed software allows for automated and high-throughput compositional analysis on DIA-MS-based glycomics spectra. As illustrated in **Figure S10A**, pre-filtering of spectra information is first performed using setting parameters, including the mass tolerance, intensity threshold, and maximum charge range. Next, the software uses an input glycan library containing glycan composition and their identified fragments to identify the glycans from both MS1 and MS2 levels. Finally, the peak of glycans can be allocated by realigning the information from MS1 and MS2, and the aligned tandem-MS spectra are saved in mgf format for further annotation. Moreover, the “add-on mass” feature allows analysis of modified glycans in samples, such as labeled, derivatized, and reduced glycans.

The models for removing false-positive results from glycomics data are hampered by the complicated non-linear glycan structures. Hong and co-workers introduced a *p*-value scoring system in GlycoDeNovo2 software.^21^ Liu *et al.* also developed GlycoNote with the random mass shift algorithm to filter false positives.^22^ For more confident glycan identification from GlycanDIA data, we optimized parameters adopted from GlycoNote and embedded them in an optional decoy mode of the software. Two parameters, coverage of intensity (Cov. Int.) and coverage of sequence (Cov. Seq.) were calculated to represent the relative abundance of matched fragments and fragmentation efficacy. For each aligned tandem MS, 100 decoy sets were generated, and the average distribution of Cov. Int. and Cov. Seq. was estimated by iteratively searching against the decoy database (**Figure S10B**). As shown in **Figure S11**, we reconstructed spectra of glycan Hex(5)HexNAc(5)Neu5Ac(1) from GlycanDIA data using the software and observed an excellent match to the reference spectra from DDA analysis. By manually checking all the identified glycans after filtering, we confirmed the performance of the software, and the detailed instructions and results about the software are described in **Supplementary Information 2**.

### GlycanDIA empowers sensitivity and accuracy for glycomic analysis

To illustrate the performance of the GlycanDIA workflow in characterizing glycans under the most native and intricate scenarios, we have employed the analysis to characterize *N*-glycans and HMOs in their protonated native forms and *O*-glycans in their reduced forms. First, we characterized the *N*-glycans released from a mixture of commercially available glycoprotein standards containing RNase B and human sera. As a result, the GlycanDIA approach outperformed DDA in terms of both identification power and quantification accuracy. Specifically, DIA consistently mapped over 270 different glycans (including isomers) from three replicates, while DDA only identified around 200 glycans (**Figure 2C**). We further examined each glycan from the released standards and selected at least five most abundant interference-free transitions to sum for glycan quantification, and we found significant improvements in the limit of quantification (LOQ) and coefficients of variation (CVs). As shown in **Figure S12**, the glycan peak can be barely quantified after a 100-fold dilution of the standard using the DDA method on the nanoLC-Time-of-Flight (ToF) system, while decent linearities were achieved even after a 1000-fold dilution for most glycans using GlycanDIA (**Figure S13**). In addition, the CVs of the technical replicates were less than 5% on average when using the more selective fragment ion measurements (**Figure 2D**). Overall, the results suggest that the GlycanDIA workflow can increase the number of glycan identification by 25% over the conventional DDA method while generating more sensitive and consistent quantitative results. Importantly, all the glycans annotated and quantified from DDA analyses were able to be distinguished using GlycanDIA (**Figure 2E**).

To demonstrate the advantages of GlycanDIA for glycomic analysis, we spiked three standard heavy isotope-labeled N-glycans at the attomolar level into the N-glycan pool released from RNase B and human sera. We then analyzed the sample with ten continuous runs using the DDA or GlycanDIA method. Notably, the GlycanDIA method provides more consistent glycan identification and stable quantitive results, while the conventional DDA method displayed limitations in such low concentrations of glycans in terms of randomly missing identification and a high level of quantification variability (**Figure 2F** and **G**).

Besides *N*-glycans, analysis of other types of glycans, particularly HMOs and O-glycans, is also critical for investigating different biological processes.^7^ Therefore, we examined the generalizability of GlycanDIA for characterizing HMOs and *O*-glycans. To adapt the method, we merely changed the mass window range to 300-1600 *m/z* because HMOs and *O*-glycans are relatively smaller compared to N-glycans. As shown in **Figure 2H**, 15 different HMO standards can all be monitored at the femtomole level. We then monitored the HMOs from the human milk sample; the signal of 163.06 *m/z* and 204.08 *m/z* at the MS2 level shows the general level of hexose and *N*-acetylhexosamine-containing HMOs (**Figure S14A**). As a result, more than 70 HMOs can be confidently quantified (**Figure S14B**). We also investigated the GlycanDIA on *O*-glycans released from bovine mucin. As shown in **Figure S15A** and **B**, more than 35 *O*-glycans were identified, and the tandem MS signal of reduced *N*-acetylhexosamine (224.11 m/z) aligned the *O*-glycan profiles, which confirmed *O*-glycans were generated from *O*-GalNAc-linked mucins. Surprisingly, the sulfated *O*-glycans normally have a lower response in the positive mode compared to the negative mode;^23^ however, several sulfated *O*-glycans can still be identified from GlycanDIA, which suggests the sensitivity of the method. Overall, the results showed the exceptional sensitivity and accuracy of GlycanDIA for glycomic analysis.

### Isomers and modified glycans can be determined using GlycanDIA

Glycan isomers are important features for various biological events. The combination of different monosaccharides often generates similar masses for some glycans. For example, the mass of one fucose adding one *N*-glycolylneuraminic acid (Neu5Gc) equals one hexose adding one Neu5Ac (453.14 Da), and the mass of two fucoses (292.11 Da) is similar to a single Neu5Ac (291.09 Da). Therefore, tandem MS information is needed as a layer of evidence for obtaining monosaccharide composition information and differentiating the glycan isomers. However, the data-dependent feature of DDA limited the glycomic analysis in this aspect, while GlycanDIA can overcome this by generating the product information for all the precursors. To validate the performance of GlycanDIA for distinguishing glycan isomers, we first checked the two common isomeric HMOs, 2’-fucosyllactose (2’FL) and 3-fucosyllactose (3FL), from the HMO standard pool. As shown in **Figure 2I**, they can be distinguished based on the relative abundance of lactose fragments (325.11 and 343.12 *m/z*). Indeed, 325.11 *m/z* was used to monitor 3FL during the targeted analysis.^24^ For *O*-glycan isomers from the collisional-based dissociation, Xu *et al.* previously annotated *O*-glycan Hex(2)HexNAc(2)Fuc(1)Neu5Ac(1) isomers based on their signature ions generated from the different positions of fucose and sialic acid.^25^ We examined whether this information could be resolved in GlycanDIA and found a series of aligned fragments that determined its structure, especially the signature ion 657.23 *m/z* (**Figure S15C** and **D**). To further evaluate the performance of GlycanDIA for distinguishing *N*-glycan isomers, we selected the fetal bovine serum sample containing both Neu5Ac and Neu5Gc and released the *N*-glycans from the protein. As shown in **Figure S16A**, six peaks corresponding to three major glycan groups were found after extracting the 1120.90 *m/z* ion at the MS1 level. Three different *N*-glycan compositions, Hex(6)HexNAc(4)Fuc(1)Neu5Ac(1) (6411),

Hex(5)HexNAc(4)Fuc(0)Neu5Ac(1)Neu5Gc(1) (54011), and Hex(5)HexNAc(4)Fuc(2)Neu5Ac(0)Neu5Gc(1) (54201) can have a same mass at 1120.90 *m/z*. The monoisotopic masses of 6411 and 54201 are exactly the same, but 54011 differs by around 0.5 at charge state 2. After looking into the MS1 spectra at different times, the monoisotopic peaks 1120.91, 1120.91, and 1290.39 *m/z* were identified at 29, 35, and 39 minutes, respectively. The MS2 spectrum at 39 minutes showed the diagnostic peaks of both Neu5Ac and Neu5Gc, suggesting the glycan with 54011 composition (**Figure S16B**). On the other hand, the peaks around 29 and 35 minutes contained Neu5Ac fragments, while they did not contain any Neu5Gc ions, indicating the peaks belong to the glycan 6411. Furthermore, the two 6411 isomer structures can be proposed based on their fragments and the ratio between the fragmented antenna with and without (*e.g.*, 512.20 and 657.24 *m/z*, respectively) informed the peak at 29 minutes is the 6411 glycan with antenna fucose and the other peak (at 35 minutes) is more likely to be the glycan with core fucose structure instead. It should be noted that the linkage specification is still missing in GlycanDIA because HCD cannot reveal the general linkage information. Interestingly, Pett *et al.* showed the assignment of α2,3-/α2,6-linked sialic acid from glycoproteomic analysis.^26^ To investigate if GlycanDIA data can disclose the α2,3- and α2,6-sialylated glycan isomers, we examined the *N*-glycan isomers, Hex(5)HexNAc(4)Neu5Ac(1) (5401), that have been well-characterized using PGC (**Figure S17A**).^27^ Remarkably, by normalizing the intensity of fragments to the general 366.14 *m/z* fragment, the relative abundance of several fragment ions displayed different preferences in our dataset. Similar to the observation from glycoproteomic results, α2,3-linked glycans generated more abundant product ions at 292.10 and 274.09 *m/z* compared to α2,6-sialylated glycans (**Figure S17B**). Besides, when sialic acid is located at the 3’ arm, it produces generic higher antenna ions (such as 1055.37, 893.32, 749.28, and 690.24 *m/z*) compared to 6’ arm-sialic acid. Surely, a more general inspection of sialylated N-glycans is required for further validation, while these results suggest that GlycanDIA can generate useful information for determining glycan isomers.

Similar to proteins, glycans can be modified, for example, by acetylation, sulfation, and phosphorylation; however, due to the low abundance of these modifications, a specific enrichment method is normally required for their mapping.^28^ Data generated from GlycanDIA workflow theoretically contains the PTM information because the DIA-based method can fragment all the ions within a selection window. To investigate if the modified glycans can be directly identified from the unenriched samples using GlycanDIA, we examined the MS2 ions at 699.24 *m/z*, which corresponds to the signature ions produced from the acetylated sialic acid. As shown in **Figure S18**, various fragment peaks of acetylated sialylated glycans were identified from the data, and one of the spectrum corresponding to Hex(5)HexNAc(4)SiaAc(1) glycan was shown as the example. Notably, the level of sialic acid acetylation is three orders of magnitude less than the unmodified sialic acid based on the ratio between their signature ions, and such low abundant modification can barely be identified using the traditional DDA-based method. These results emphasize the significant advantage of GlycanDIA and the potential to acquire missing information when the dataset is re-interpreted later.

### Revealing the landscape of N-glycans from RNA extracts

The recent discovery of RNA glycosylation has allowed for the mapping of RNA glycan profiles. However, mass spectrometry has yet to be successfully utilized for RNA glycan profiling due to the low level of glycosylation on RNAs (approximately 20 pmol per μg of total RNA).^1^ The GlycanDIA workflow has the substantial advantage of accurately identifying and quantifying low-abundance glycans. Therefore, we applied GlycanDIA to characterize the *N*-glycans released from RNA extracts (**Figure 3A**). To ensure the purity of the RNA, we treated the crude RNAs from TRIzol extraction with DNase and protease, and no proteins could be visualized from the SDS-PAGE gel (**Figure S19A**). This is important to rule out cross-contamination from any glycoproteins.

**Figure 3.**
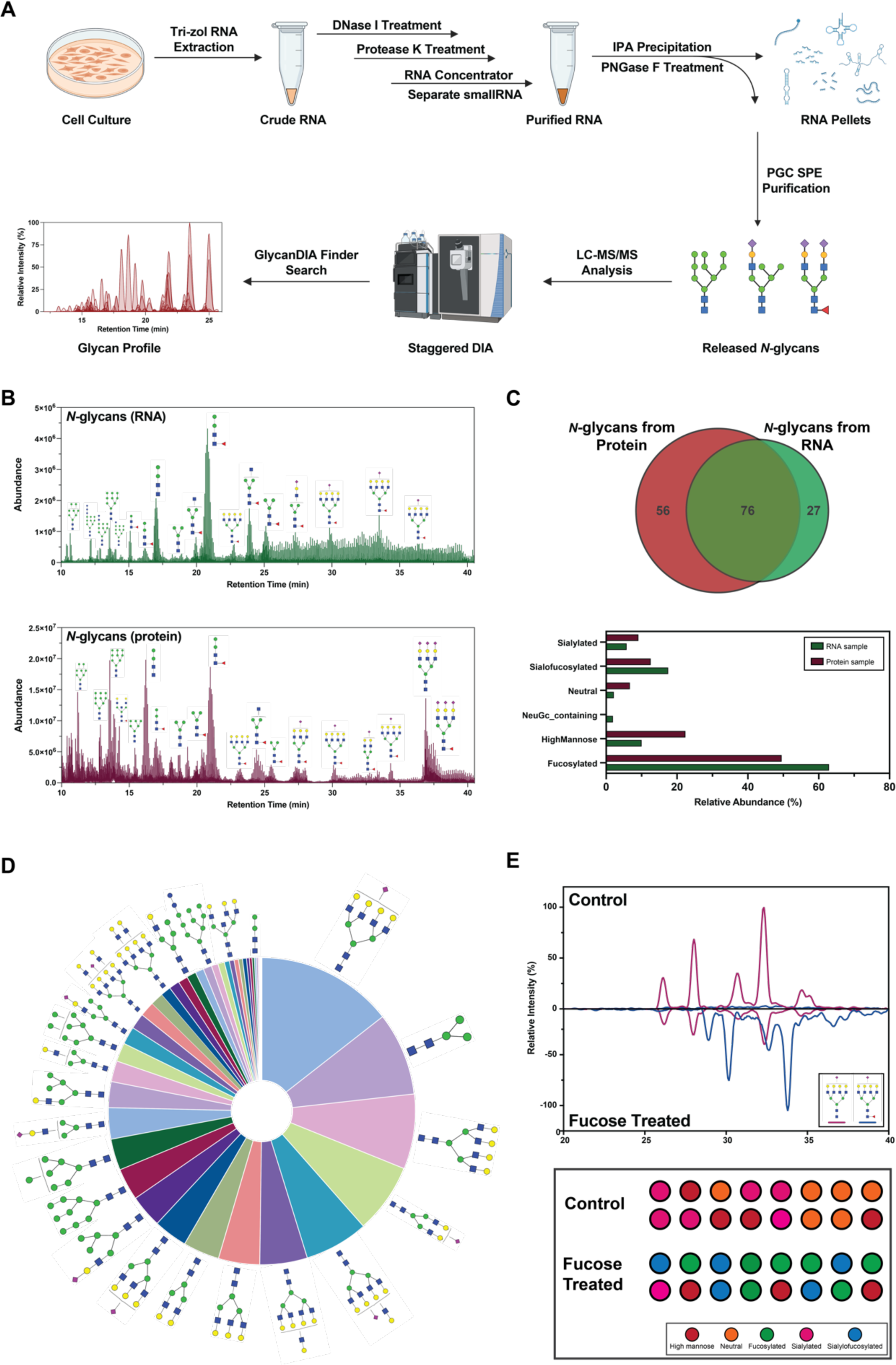
GlycanDIA provides new insights into the glycomic analysis. **A.** RNAs were extracted from cells using Trizol reagents, and small RNAs were separated using an RNA concentrator after the treatment of DNase and protease. The extracted RNAs were subject to the PNGaseF treatment to release *N*-glycans. Glycans were purified using a PGC SPE plate, analyzed using the DIA-based high-resolution mass spectrometry, and identified using GlycanDIA Finder. **B.** The results of *N*-glycans in HEK 293T RNA (top) and protein (bottom) samples showed different glycomic profiles. **C.** RNA and protein *N*-glycan profiles differed in relative abundances and structures. **D.** Top 16 RNA *N*-glycans identified from HCT116 cell lines. The fucosylated glycans were not identified due to the GMD protein mutation. **E.** *N*-glycans, such as the Hex(7)HexNAc(6)Sia(1), were converted into glycans containing fucose after the external fucose feeding.

We first monitored the glycans released from either RNAs or glycoproteins in HeLa and HEK 293T cells. After fractionating the total RNA into large and small RNAs based on size (**Figure S19B**), the signature glycan signals were observed from the small RNA fraction, while no significant glycans were identified from the large RNA fraction (**Figure S20**). As shown in **Figure 3B**, the *N*-glycans released from the HEK 293T RNA extracts showed an overall distinct profile compared to those from the glycoproteins, whereas RNA extracts were dominated by fucosylated and sialofucosylated (containing both sialic acid and fucose) glycans. Similarly, the *N*-glycan profile of RNA from HeLa cells showed more abundant sialofucosylated glycans (**Figure S21**). Over 100 glycans were identified and quantified from protein and RNA extracts, and the overlap of the identified glycans between the two populations was more than 70% (76 out of 103). The identified glycans from the glycoproteins were similar to the results from previous studies,^1^ however the GlycanDIA method used only 5 μg of small RNA (5-fold less than used in the initial study) and resulted in deeper and more robust identification of the *N*-glycans. Remarkably, although a large overlap between the two fractions was observed in different types of glycans, the abundance of individual glycoforms and subtypes varied significantly (**Figure 3C**). A potential explanation is that different glycosyltransferases and glycosidases may exhibit distinct preferences toward their proteins or RNA substrates.

We then investigated the glycans on RNAs in the human colorectal carcinoma cell line, HCT116. This cell line is a glycosylation model that lacks fucosylation due to the mutation of the GDP-mannose-4,6-dehydratase (GMD) enzyme which blocks the fucose *De Novo* synthesis.^29^ The fucosylation of the cell line’s glycoproteins can be rescued by adding external fucose through the *Salvage* pathway. Currently, this pathway has not been investigated for the fucosylation of the RNAs. As shown in **Figure 3D**, we detected zero fucosylated glycans from the HCT116 RNA extracts, and the most abundant glycan was a sialylated glycan, Hex(7)HexNAc(6)Fuc(0)Sia(1) (7601). After treating the cells with 100 mM of fucose, the glycan profile was reprogrammed towards fucosylated and sialofucosylated glycan subtypes. For example, over 90% of the sialylated glycan 7601 was converted to the sialofucosylated glycan, Hex(7)HexNAc(6)Fuc(1)Sia(1) (7611) (**Figure 3E**). This result reveals that the *N*-glycans on HCT116 RNAs can also be modified through the canonical biosynthesis enzymes in carbohydrate metabolism.

Lastly, we applied GlycanDIA to unravel the profile of glycans on RNA from different mouse tissues. As a result, more than 200 *N*-glycans in total were identified from three replicates of five different tissues (**Figure 4A**), and the landscape of glycans from tissue glycoRNAs was different from the profiles of glycoprotein samples.^30, 31^ In addition, the relative abundance of different N-glycan subtypes was also distinct (**Figure 4B**). For example, heart tissue exhibited relatively more abundant high mannose type glycans (30%), and fucosylated glycans were predominant with over 75% of total glycans identified from the brain. Most tissues showed higher ratios of sialic acid-containing glycans, being consistent with the total degree of sialylation we assessed using the RNA periodate oxidation and aldehyde labeling (rPAL) method (**Figure S22**).^32^ Furthermore, there is a variation in the most prevalent glycoforms among different tissues. For example, the glycans consisting of tetraantennary Hex(7)HexNAc(6) and Hex(7)HexNAc(7) structures were found to be highly abundant in the colon, while these glycans were barely identified from other samples. Strikingly, although sialylated structures account for more than 50% of glycans in most tissues, the pattern of sialylation (Neu5Ac and Neu5Gc) varied across different tissues (**Figure 4C**). For example, the colon and the heart had relatively higher Neu5Ac, while the spleen exhibited a higher degree of Neu5Gc. These results were also confirmed with the base peak chromatogram (BPC) of Neu5Ac and Neu5Gc signature ions (**Figure S23**). Overall, the results demonstrate details about the abundance and specific distribution of *N*-glycans in different glycoRNA samples, which can be a critical feature that regulates cellular interactions and governs cellular functions.

**Figure 4.**
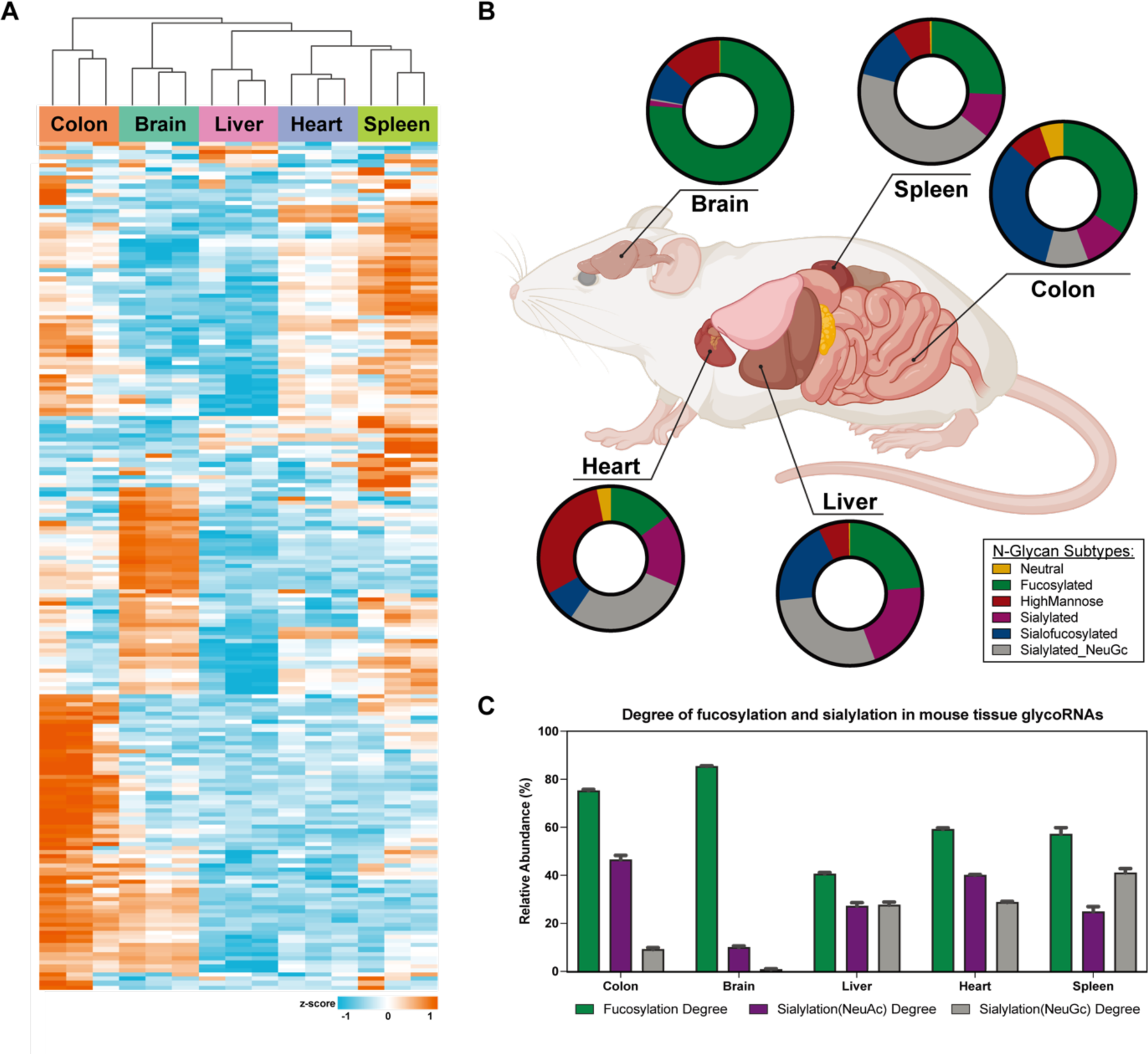
*N*-Glycan profiles of glycoRNA from mouse tissues. **A.** Various *N*-glycans in total were identified from different mouse tissues, and the heatmap showed distinct *N*-glycan profiles from the different mouse tissues. **B.** The differences in the relative abundance of *N*-glycan subtypes were also noticed in the mouse tissues. **C.** The high degree of fucosylation was generally noted in all different tissue RNAs, while the degree of sialyation and the level of Neu5Ac and Neu5Gc were tissue-specific.

## Discussion

DDA-based methods have been prevalent in current glycomic studies. However, the nature of selecting the most abundant ions in DDA often impedes the investigation of subtle glycosylation changes in complex biological samples, especially when the amount of material for analysis is limited. As our showcase in analyzing standard heavy isotope-labeled N-glycans, the detection and quantification from DDA are inconsistent at the attomolar level; while the robust identification and quantitative results from GlycanDIA demonstrate its capability to capture glycan information with lower sample demands. It is noted that the strategy of DIA can always get the best fragmentation at (or near) the apex of the compound peak, while DDA can hardly be promised to trigger at the apex, even with the system optimization or apex-trigger algorithm.^33^ This means the GlycanDIA can generally produce glycan fragmentation with good quality and higher sensitivity. For example, in our analysis of glycans on HEK 293T RNA, we identified nearly 100 glycans using only 5 μg of small RNAs, while 25 μg of HEK 293T small RNAs were used in the previous study, finding only 14 *N*-glycans. This type of increased sensitivity and depth will enable others to more easily study how cells glycosylate various template biopolymers across a wider array of biological conditions. Although the parallel reaction monitoring (PRM) method was recently introduced to glycomic analysis, providing high sensitivity and high resolution, pre-knowledge about the analytes (such as target mass and retention time) is required, and only the ions within the inclusion list can be fragmented.^34^ Meanwhile, GlycanDIA workflow owes the unique advantage that the dataset can continually be re-analyzed since all the precursors in the set mass ranges were fragmented during the data acquisition. This can provide valuable information by re-evaluating glycan modifications and discovering new glycan structures in future studies. Importantly, GlycanDIA provides a universal tool to monitor the level of glycosylation from different samples, which can be beneficial for identifying novel glycosylated molecules. For a more direct comparison, the general features of different glycomics methods are listed in **Table S4**.

Previously, the quantification of glycans was only investigated at the MS1 level since MS2-based quantification was considered to be ambiguous. This is because, unlike peptides, glycans often generate common fragments such as hexose (163.06 *m/z*) and *N*-acetylhexosamine (204.08 *m/z*) during the fragmentation event, which may be produced by multiple co-eluting glycans and hardly used for the MS2-based quantification. Meanwhile, unique glycan fragments from CID have been commonly monitored for targeted MRM-based glycomic analysis, which inspired us to develop the GlycanDIA approach.^12, 24^ The targeted method essentially is a special DIA-like approach that selects small and discrete mass windows. In this study, we established the pipeline for data analysis using GlycanDIA, which provides essential information on glycan identities and quantities. Although some glycomic databases are available, the spectra are mostly constructed under negative CID or HCD, which can not be directly applied to our current workflow using HCD in positive mode. Therefore, we first generated predicted fragments for all possible *N*-glycans using GlycoWorkbench, followed by selecting the fragments that can represent the corresponding glycans based on the analysis of DDA data using Skyline. We showed that MS2-based quantification is feasible in glycomic analysis by GlycanDIA after meticulously optimizing the isolation window and picking multiple co-varying transitions. Quantification using the MS2 signals is generally more sensitive because the MS1 signals of a glycan are more likely to have interference from another analyte in the sample with a smaller mass, while the selective fragment ion measurements in MS2-based quantification approach can be less prone to noise interference. Notably, due to the potential gas phase rearrangement of fucose, having a good separation of glycans and carefully choosing the product ions that represent the parent glycans during the GlycanDIA is crucial.^35, 36^

Glycomics in biological analysis is more often focused on elucidating the glycan composition information, and GlycanDIA can provide superior discovery of such information. Although we have distinguished example glycan isomers from HMO, *O*-glycan, and *N*-glycan samples, the current GlycanDIA is still limited in detailing the linkage between each monosaccharide residue and providing better utility for linkage determination due to the major glycosidic bond cleavage during HCD fragmentation. The linkage of sialic acid and fucose can be readily monitored by comparing the glycan peaks before and after the treatment of sialidases and fucosidases.^25^ However, the integral glycan structure characterization requires other types of fragmentations and applications of the MS^n^ strategy. For example, Wei *et al.* reported the investigation of the glycan linkages with remarkable details using MS^3^ and electron-based fragmentations.^37^ In addition, indexed time correction (iRT) for DIA analysis is a critical parameter for the analysis between different samples and batches to increase confidence in identification. Ashwood *et al.* demonstrated the use of glucose oligomers for normalizing retention values during glycomic analysis on a PGC column.^38^ Therefore, the improvement of GlycanDIA in the future can be focused on elucidating glycan structural information using other fragmentation techniques and matching their retention time with iRT calibration, which can ideally facilitate the detailed identification of glycans using GlycanDIA. At the current stage, we believe GlycanDIA enables a more comprehensive glycomic analysis compared to the DDA-based methods, offering a unique aspect that is rarely explored in glycomics.

## Conclusions

Glycosylation is a prevalent feature of disease progression, reported in cases of cancer, diabetes, and Alzheimer’s disease. The different glycosylation significantly influences the biological activity of proteins via alternating glycoprotein structures and potentiating the binding of receptors. We reported an integrative workflow, GlycanDIA, to characterize the glycans using the cutting-edge DIA strategy. Our results exemplify the workflow’s advantages in improved detection and quantification to provide a comprehensive view of different cellular glycosylation, including *N*-glycans, *O*-glycans, and HMOs. Meanwhile, information about glycan isomers and low-abundant glycan modifications can also be revealed. Such glycomic information can be used to guide the design of therapeutics with coordinated binding to site-specific glycosylated targets.

## Methods Method Details

### Cell Membrane Extraction

The procedures for cell membrane extraction were described previously.^39^ Briefly, cell pellets were lysed on ice with five alternating on and off pulses in 5 and 10-second intervals using a probe sonicator (Fisher Scientific, NH). Nuclear and cellular debris was removed by centrifugation at 2000 × *g* for 10 min at 4 °C. The supernatants were then centrifuged at 200,000 × *g* for 45 min at 4 °C using Optima Max-XP Ultracentrifuge (Beckman, IN) to extract the plasma membrane. The pellets of the cell membrane were washed with 0.2 M Na2CO3 solution and water, respectively.

### RNA Extraction

RNA extraction and processing took place as described.^32^ Specifically, samples were thawed after homogenization and denaturing was further encouraged by placing the samples at 50 °C and shaking for 5 minutes. To phase separate the RNA, 0.4X volumes of water was added, vortexed, let to stand for 5 minutes at 25 °C and lastly spun at 12,000x *g* at 4 °C for 15 minutes. The aqueous phase was transferred to clean tubes and 1.1X volumes of isopropanol was added. The RNA is then purified over a Zymo-II column (Zymo Research, CA). First, 350 μL of pure water was added to each column and spun at 10,000x *g* for 30 seconds, and the flowthrough was discarded. Next, precipitated RNA from the RNAzol RT extraction (or binding buffer precipitated RNA, below) is added to the columns, spun at 10,000x *g* for 10-20 seconds, and the flowthrough is discarded. This step is repeated until all the precipitated RNA is passed over the column once. Next, the column is washed three times total: once using 400 μL RNA Prep Buffer (3 M GuHCl in 80% EtOH), twice with 400 μL 80% ethanol. The first two spins are at 10,000x *g* for 20 seconds, the last for 30 sec. The RNA is then treated with Proteinase K on the column. Proteinase K is diluted 1:19 in water and 50 μL added directly to the Zymo-II column matrix, and then allowed to incubate on the column at 37 °C for 45 minutes. The column top is sealed with either a cap or parafilm to avoid evaporation. After the digestion, the columns are brought to room temperature for 5 minutes; lowering the temperature is important before proceeding. Next, eluted RNA is spun out into fresh tubes and a second 50 μL elution with water is performed. To the eluate, 1.5 μg of the mucinase StcE is added for every 50 μL of eluted RNA, and placed at 37 °C for 30 minutes to digest. The RNA is then cleaned up again using a Zymo-II column. Here, 2X RNA Binding Buffer (Zymo Research, CA) was added and vortexed for 10 seconds, and then 2X (samples + buffer) of 100% ethanol was added and vortexed for 10 sec. This is then bound to the column, cleaned up as described above, and eluted twice with 50 μL water. The final enzymatically digested RNA is quantified using a Nanodrop. To then isolate small RNAs, we followed the “Purification of Small and Large RNAs into Separate Fractions” protocol in the RNA Clean & Concentrator 5 (Zymo Research, CA) protocol exactly as described.

### rPAL labeling from mouse tissues

we followed the procedure as described.^32^ We started by lyophilizing 1 μL of enzymatically treated small RNA from above. First, we prepared blocking buffer: for each reaction we mixed 1 μL 16 mM mPEG3-Ald (BroadPharm, CA), 15 μL 1 M MgSO2 and 12 μL 1 M NH4OAc pH 5 (with HCl). 28 μL of the blocking buffer is added to the lyophilized RNA, mixed completely by vortexing, and then incubated for 45 minutes at 35 °C to block. The samples are briefly allowed to cool to room temperature (2-3 minutes), then working quickly, 1 μL 30 mM aminooxy-biotin (Cayman Chemicals, MI) is added first, then 2 μL of 7.5 mM NaIO4 (periodate, stock made in water) is added. The periodate is allowed to perform oxidation for exactly 10 minutes at room temperature in the dark. The periodate is then quenched by adding 3 μL of 22 mM sodium sulfite. The quenching reaction is allowed to proceed for 5 minutes at 25 °C. Both the sodium periodate and sodium sulfite stocks were made fresh within 20 minutes of use. Next, the reactions are moved back to the 35 °C heat block, and the ligation reaction is allowed to occur for 90 minutes. The reaction is then cleaned up using a Zymo-I column. 19 μL of water is added in order to bring the reaction volume to 50 μL, and the Zymo protocol is followed as per the above details. RNA was eluted from the column using 2x 6.2 μL water (final volume approximately 12 μL).

### RNA blotting from mouse tissues

In order to visualize the periodate labeled RNA, it is run on a denaturing agarose gel, transferred to a nitrocellulose (NC) membrane, and stained with streptavidin as described.^32^ RNA is combined with 12 μL of Gel Loading Buffer II (GLBII, 95% formamide, 18 mM EDTA, 0.025% SDS) with a final concentration of 2X SybrGold and denatured at 55 °C for 10 minutes. It is important to not use GLBII with dyes. Immediately after this incubation, the RNA is placed on ice for at least 2 minutes. The samples are then loaded into a 1% agarose, 0.75% formaldehyde, 1.5X MOPS Buffer (Lonza Bioscience, NC) denaturing gel. Precise and consistent pouring of these gels is critical to ensure a similar thickness of the gel for accurate transfer conditions; we aim for approximately 1 cm thick of solidified gel. RNA is electrophoresed in 1x MOPS at 115V for between 34 or 45 minutes, depending on the length of the gel. Subsequently, the RNA is visualized on a UV gel imager, and excess gel is cut away; leaving ∼0.75 cm of gel around the outer edges of sample lanes will improve transfer accuracy. The RNA is transferred using transfer buffer (3 M NaCl solution at pH 1, with HCl) to a NC membrane for 90 minutes at 25 °C. Post transfer, the membrane is rinsed in 1x PBS and dried on Whatman Paper (GE Healthcare, MA). Dried membranes are rehydrated in Intercept Protein-Free Blocking Buffer, TBS (Li-Cor Biosciences, NE), for 30 minutes at 25 °C. After the blocking, the membranes are stained using Streptavidin-IR800 (Li-Cor Biosciences, NE) diluted 1:5,000 in Intercept blocking buffer for 30 minutes at 25 °C. Excess Streptavidin-IR800 was washed from the membranes using three washes with 0.1% Tween-20 in 1x PBS for 3 minutes each at 25 °C. The membranes were then briefly rinsed with PBS to remove the Tween-20 before scanning. Membranes were scanned on a Li-Cor Odyssey CLx scanner (Li-Cor Biosciences, NE).

### *N*-Glycan Sample Preparation

The cell membrane or RNAs were resuspended with 200 µL of 100 mM HEPES buffer, and the mixture was heated using a thermomixer at 100 °C for 3 min. The cleavage of *N*-glycans was performed by adding 4 µL of PNGase F (500,000 units/mL), followed by incubation in a 37 °C thermomixer overnight. The proteins and RNAs were precipitated by adding ethanol and incubating at −80 °C for 2 h, followed by centrifuging at 20,000 × *g* for 15 min at 4 °C. The supernatant containing *N*-glycans was purified using the porous graphitic carbon (PGC) SPE plate (Thermo Scientific, MA). *N*-glycans were washed with 0.1% (*v*/*v*) TFA in water and were eluted with 60% (*v*/*v*) ACN and 0.1% (*v*/*v*) TFA in water. The purified glycans were dried using the SpeedVac system (Thermo Scientific, MA) and reconstituted in water for LC-MS/MS analysis.

### *O*-Glycan Sample Preparation

The bovine mucin was mixed with 10 μL of 2 M NaOH and 100 μL of 2 M NaBH4 and incubated at 45 °C for 18 h. After the reaction, 10% acetic acid was added to the sample on ice until the pH reached acidic. Followed by centrifugation at 21,000 × *g* for 20 min at 4 °C, the supernatant containing free *O*-glycans was loaded onto the PGC SPE plate and followed as per the above details for purification. The elutes were dried, reconstituted in 89% (*v*/*v*) acetonitrile with 1% (*v*/*v*) trifluoroacetic acid in water, and further purified by iSPE-HILIC cartridges (HILICON, Sweden).

### HMO Sample Preparation

The procedures for preparing HMO samples were described previously.^40^ Briefly, the human milk sample (Milk Bank at Austin) was mixed with 90 μL of water and defatted by centrifugation at 3200 × *g* for 30 min at room temperature. The aqueous layers were transferred into new plates. Then 2 volumes of ethanol were added to precipitate the proteins at −80 °C for 2 h. After 30 min of centrifugation at 21,000 × *g* for 20 min at 4 °C, the supernatant fluids containing mainly oligosaccharides were purified using the PGC SPE plate to remove lactose.

### LC-MS/MS Analysis

The glycans were analyzed using a Vanquish Neo UHPLC System (Thermo Scientific, CA) coupled with an Orbitrap Exploris 240 mass spectrometer (Thermo Scientific, CA). 2 μL of the sample was injected, and the analytes were separated on a self-packed nano PGC column (3 μm, 0.075 mm × 250 mm). In the comparison experiment, 6520 Accurate Mass Q-TOF LC/MS equipped with a PGC nano-chip (Agilent, CA) was used. A binary gradient using solvent A with 0.1% (*v*/*v*) FA in water and solvent B with 0.1% (*v*/*v*) FA in ACN was applied to separate *N*-glycans at a 300 nL/min flow rate. The detailed parameters for MS setup are available in **Table S1**.

### Data Analysis

For DDA results, the glycans were identified and quantified using GlycoNote and Agilent MassHunter Qualitative Analysis software (v.B08).^22^ For DIA results, GlycoWorkbench (v2.1) was used to predict the glycan fragments, ProteoWizard MSConvert (v3.0) was used for demultiplexing the spectrum, and Thermo Scientific FreeStyle (v1.8) and Skyline (v21.0) software were used for viewing MS1 and MS2 Information.^41–43^ GlycanDIA Finder was used to identify and quantify the glycans. The legend for saccharide units and annotations can be found in **Table S3**.

### Detemrination of glycan abundance

The glycan subtypes were annotated as high mannose, neutral, fucosylated, sialylated, sialofucosylated, and Neu5Gc-containing based on the glycan compositions. To calculate the relative abundance, we employed the label-free approach using the intensity (height or area under the peak) of the specific glycan or subgroup glycans and normalized it to the total abundance of all the identified glycans.

## Data and code availability

The supplementary information includes all data generated or analyzed during this study. The raw mass spectrometry data have been deposited in the Mass Spectrometry Interactive Virtual Environment (MassIVE) database under accession number MSV000093677. The source code can be accessed at https://github.com/ChenfengZhao/GlycanDIAFinder. Any additional information in this paper should be directed to and will be fulfilled by the lead contacts, C.B.L., R.A.F., and B.A.G.

## Author contributions

Y.X. conceived the project, designed and performed the experiments, analyzed data, produced figures, and drafted the manuscript. X.L., S.C., Z.L., and F.M.R. performed the MS analysis and edited the manuscript. C.Z. and S.W. wrote the code and developed the search engine. performed the method setup. B.M.G. and R.A.F. performed the glycoRNA experiments and edited the manuscript. C.B.L. planned the overall experimental project and edited the manuscript. B.A.G. supervised the overall experimental project and edited the manuscript.

## Acknowledgments

The authors appreciate the Skyline and GlycoWorkbench development teams for creating this complementary software for the community. The authors also thank Maurice Wong (University of California, Davis) for the active discussion and Aaron Stacy and Yasmine Bouchibti (University of California, Davis) for preparing HMO samples. Research reported in this publication was supported by grants from the Burroughs Wellcome Fund Career Award for Medical Scientists (R.A.F.), the Sontag Foundation Distinguished Scientist Award (R.A.F.), the Rita Allen Foundation (R.A.F.), National Institutes of Health GM049077 (C.B.L.), AG062240 (C.B.L.), AI118891 (B.A.G.), HD106051 (B.A.G.), and CA196539 (B.A.G.).

## Supplementary Information 1

### Materials

Sodium carbonate (Na2CO3), guanidinium chloride (GuHCl), sodium hydroxide (NaOH), sodium borohydride (NaBH4), ammonium acetate (NH4OAc), sodium periodate (NaIO4), Magnesium sulfate (MgSO2), sodium sulfite, Tween-20, and mucinase StcE were purchased from Sigma-Aldrich, MO. 4-(2-hydroxyethyl)-1-piperazineethanesulfonic acid (HEPES) buffer, SybrGold, Proteinase K, acetonitrile (ACN), trifluoroacetic acid (TFA), and formic acid (FA) were purchased from Thermo Fisher Scientific, MA. The source of standard glycoproteins, *N-*glycans, and HMOs, were listed in **Table S2**.

### Animals

All mouse procedures and protocols were approved by the Animal Care and Use Committee of Boston Children’s Hospital and followed all relevant guidelines and regulations. C57BL/6 mice were crossed and bred in house. Male C57BL/6 mice (24-28 weeks old) were euthanized, and their liver, spleen, colon, heart, and brain were harvested. The organs were directly added to 1 mL of RNAzol RT (Molecular Research Center, Inc.) in 2 mL tubes containing Zirconium oxide beads (Thomas Scientific). Tissues were homogenized using a Bead mill 24 Homogenizer (Fisher Scientific); 4-8 cycles of 20 seconds on, 10 seconds off at 25°C. After homogenization was complete, samples were stored at −80°C or processed to extract RNA.

### Cell lines

Immortalized human embryonic kidney HEK 293T cells, immortalized human cervix HeLa cells, and human colorectal carcinoma HCT116 cells were obtained from American Type Culture Collection (ATCC, Manassas, VA). The cells were cultured in Dulbecco’s Modified Eagle Medium (DMEM) medium supplemented with 10% (*v*/*v*) fetal bovine serum (FBS), 1% (*v*/*v*) non-essential amino acids, and 1% (*v*/*v*) GlutaMAX. Cells were maintained in a humidified incubator at 37 °C with 5% CO2 and subcultured at 80% confluency.

## Supplementary Tables

**Table S1.**
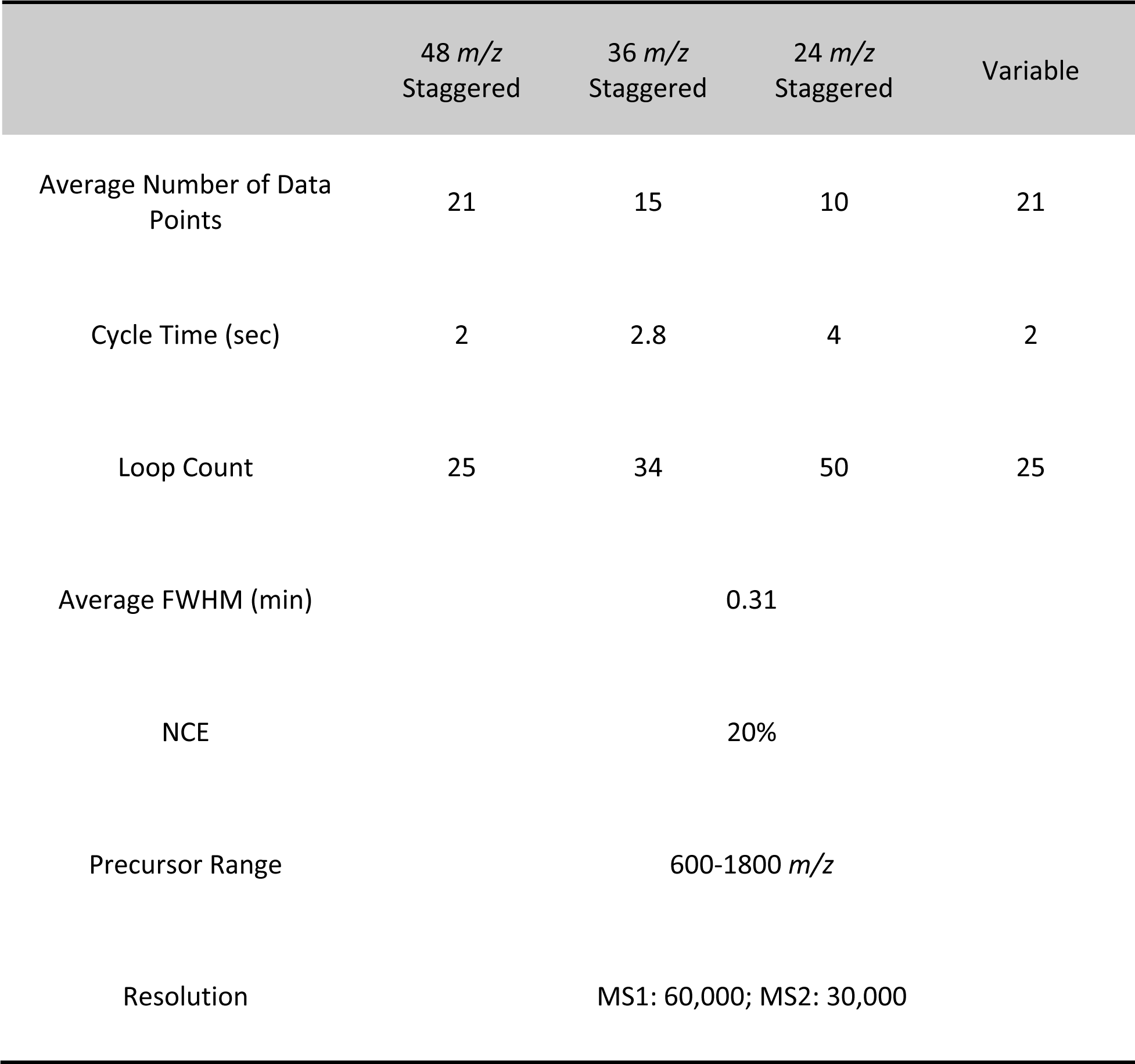
Summary of the details of LC-MS/MS parameters in GlycanDIA workflow.

**Table S2.**
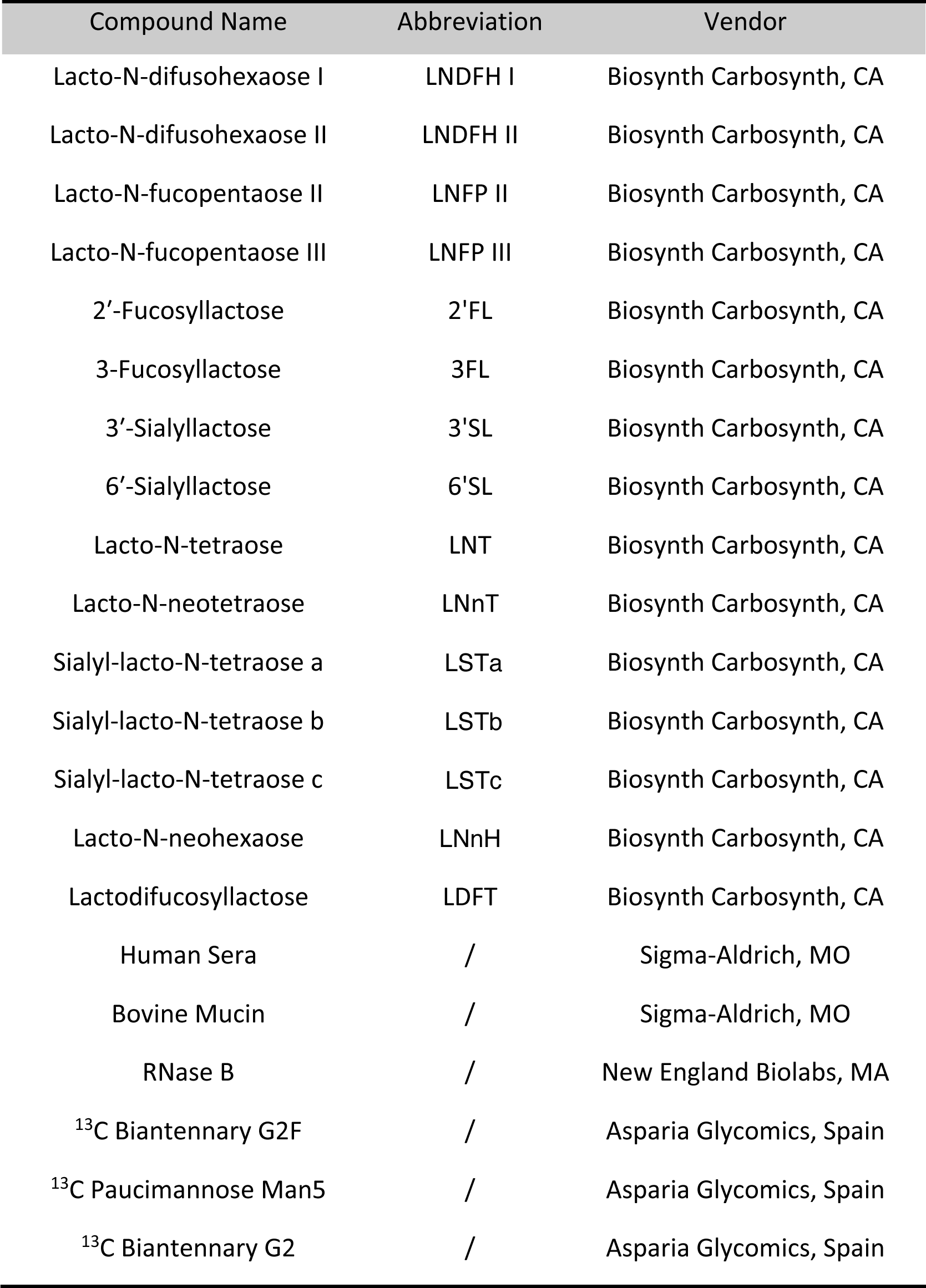
Vendor of standard HMOs, *N-*glycans, and glycoproteins.

**Table S3.**
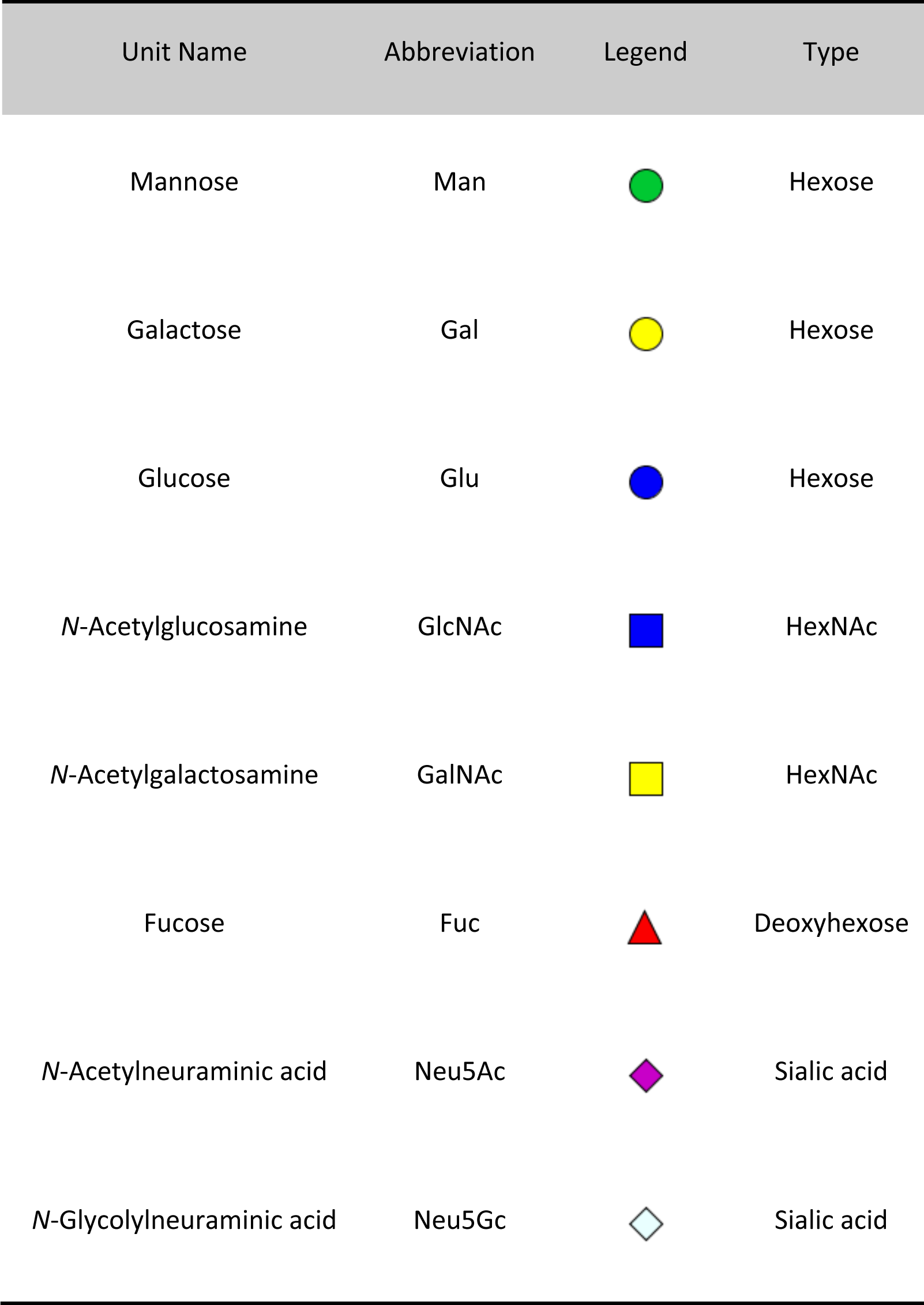
Legends and annotations of glycan units. The legend and annotation followed the rules of Symbol Nomenclature for Glycans (SNFG), and the annotation for glycans in this manuscript is as follows: Hex_HexNAc_Fuc_Neu5Ac_Neu5Gc.

**Table S4.**
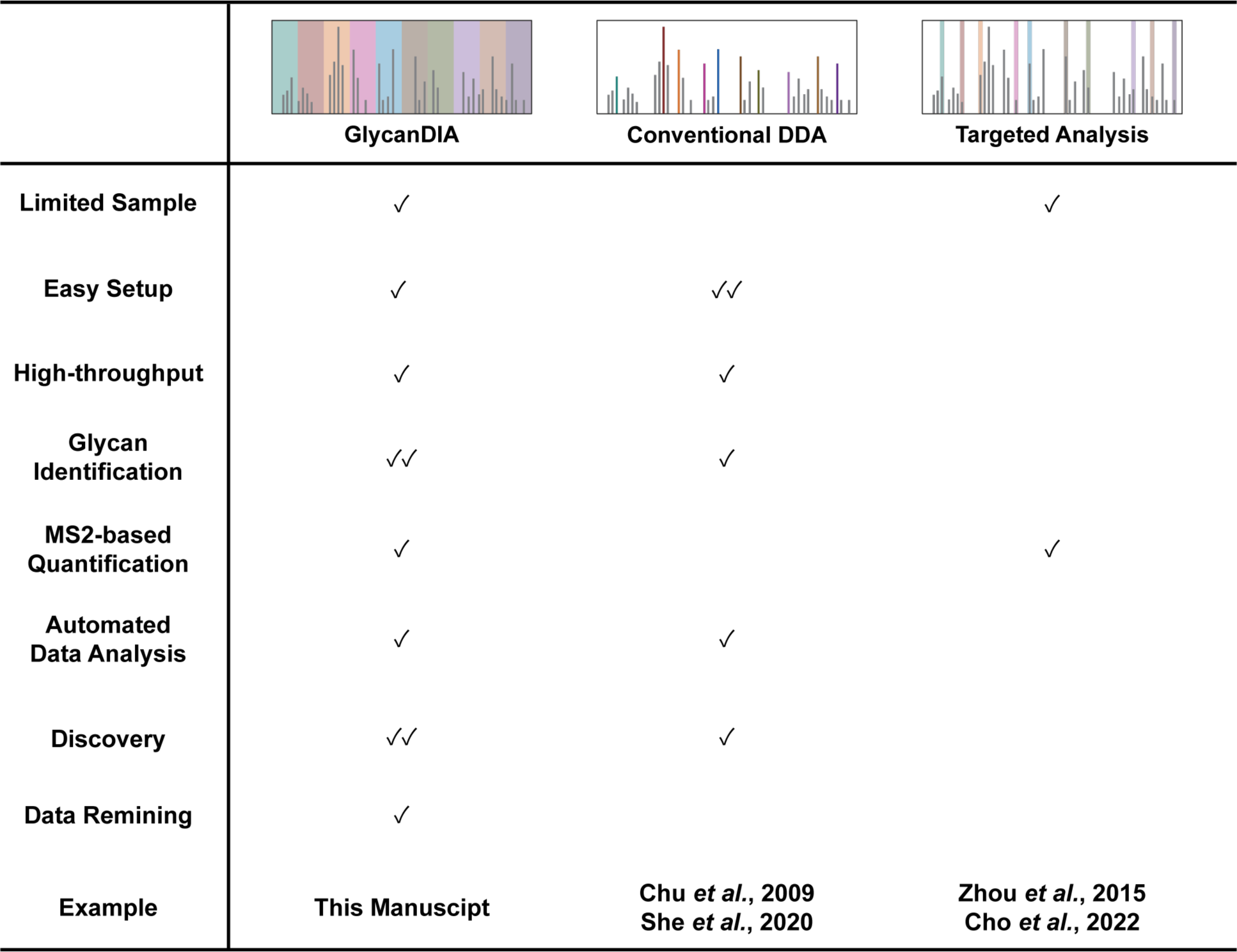
Features of MS-based methods in glycomic analysis. References of the example: Chu *et al.*, 9, 7, 1939–1951, *Proteomics* 2009; She *et al.*, 92, 20, 14038–14046, *Anal. Chem.* 2020; Zhou *et al.*, 26, 4, 596-603, *J. Am. Soc. Mass Spectrom.* 2015; Cho *et al.*, 94, 44, 15215–15222, *Anal. Chem.* 2022.

|  | **GlycanDIA** | **Conventional DDA** | **Targeted Analysis** |
| --- | --- | --- | --- |
| Limited Sample | ✓ |  | ✓ |
| Easy Setup | ✓ | ✓✓ |  |
| High-throughput | ✓ | ✓ |  |
| Glycan Identification | ✓✓ | ✓ |  |
| MS2-based Quantification | ✓ |  | ✓ |
| Automated Data Analysis | ✓ | ✓ |  |
| Discovery | ✓✓ | ✓ |  |
| Data Remining | ✓ |  |  |
| Example | This Manuscript | Chu *et al.*, 2009<br>She *et al.*, 2020 | Zhou *et al.*, 2015<br>Cho *et al.*, 2022 |

## Supplementary Figures

**Figure S1.**
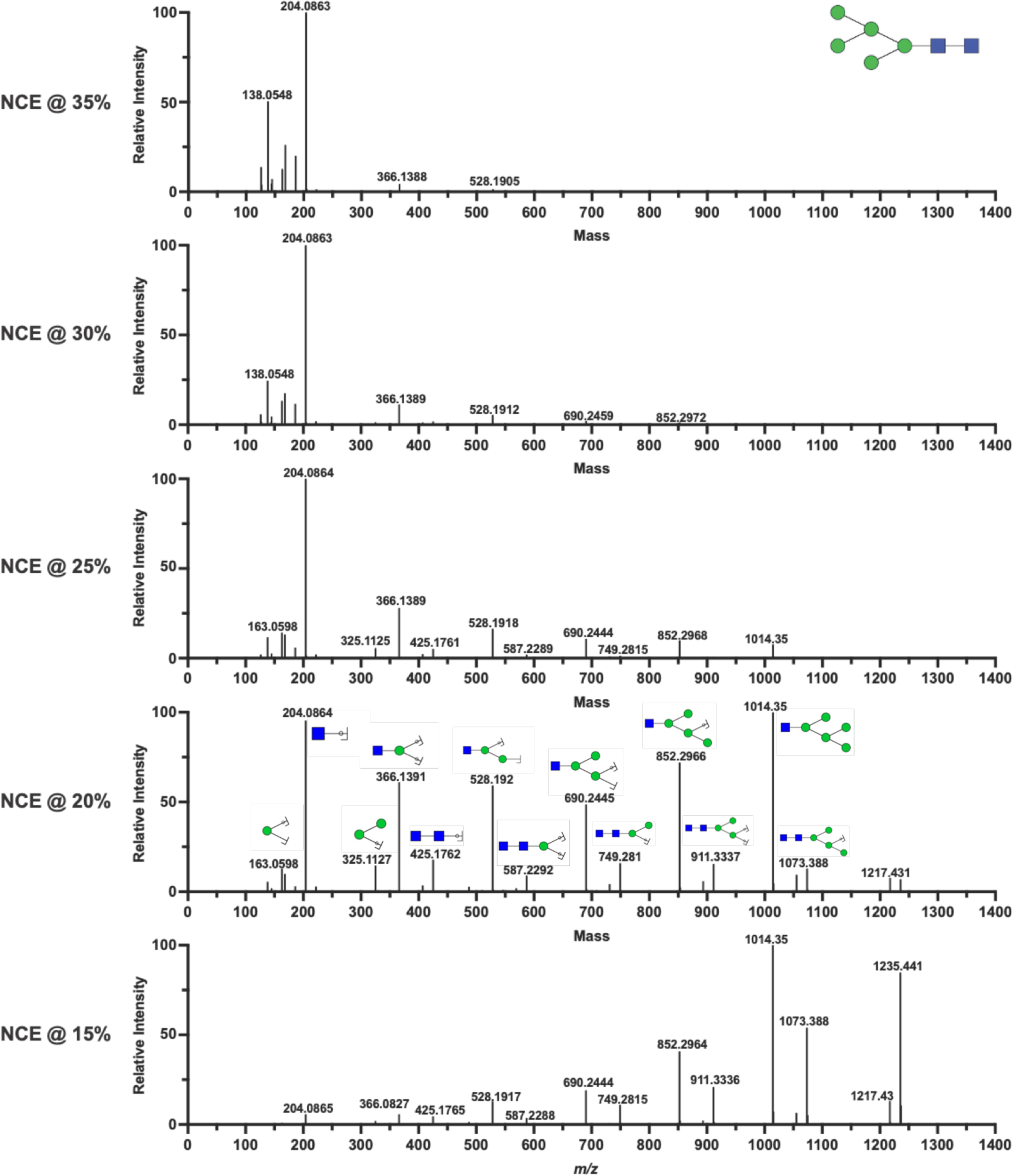
Representative tandem MS/MS spectra of high mannose glycan, Hex(5)HexNAc(2), at different fragmentation energies. The abundance of generated fragments that can be used for sequencing the example glycan increased with the NCE boost, while the sequencing information was lost after the collision energy was greater than 25% due to the over-fragmentation.

**Figure S2.**
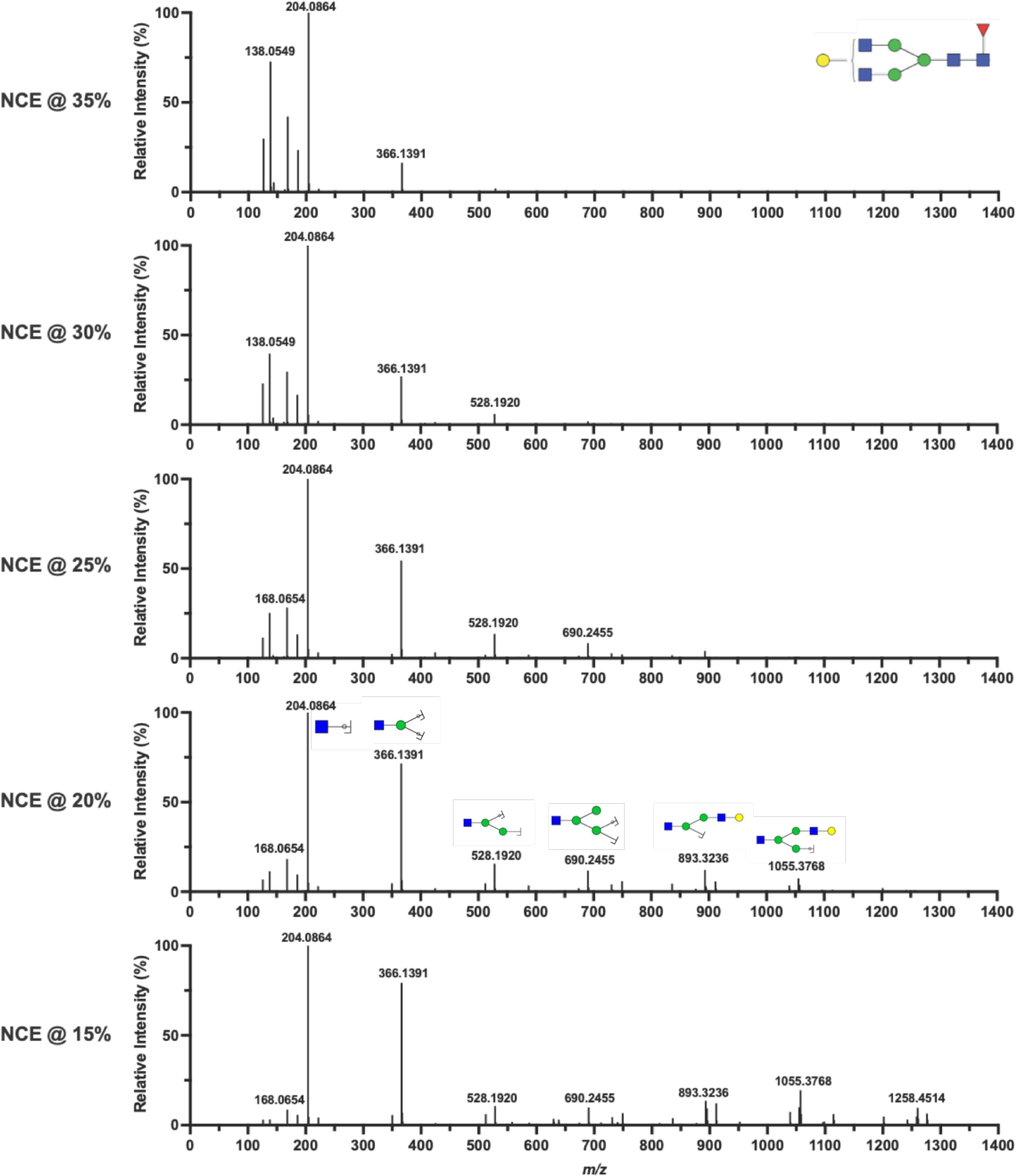
Representative tandem MS/MS spectra of fucosylated glycan, Hex(4)HexNAc(4)Fuc(1), at different fragmentation energies. The abundance of generated fragments that can be used for sequencing the example glycan increased with the NCE boost, while the sequencing information was lost after the collision energy was greater than 25% due to the over-fragmentation.

**Figure S3.**
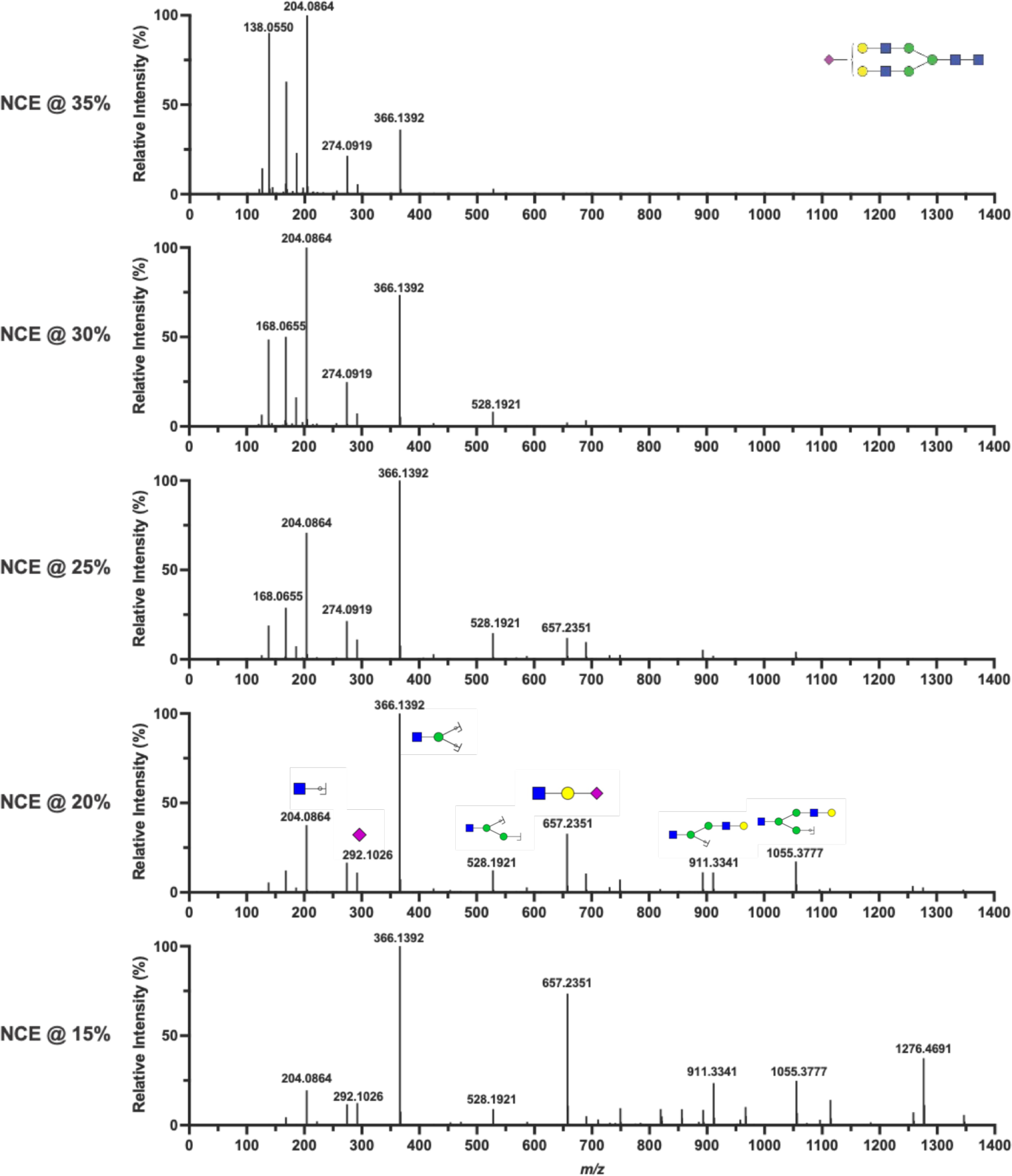
Representative tandem MS/MS spectra of sialylated glycan, Hex(5)HexNAc(4)Sia(1), at different fragmentation energies. The abundance of generated fragments that can be used for sequencing the example glycan increased with the NCE boost, while the sequencing information was lost after the collision energy was greater than 25% due to the over-fragmentation.

**Figure S4.**
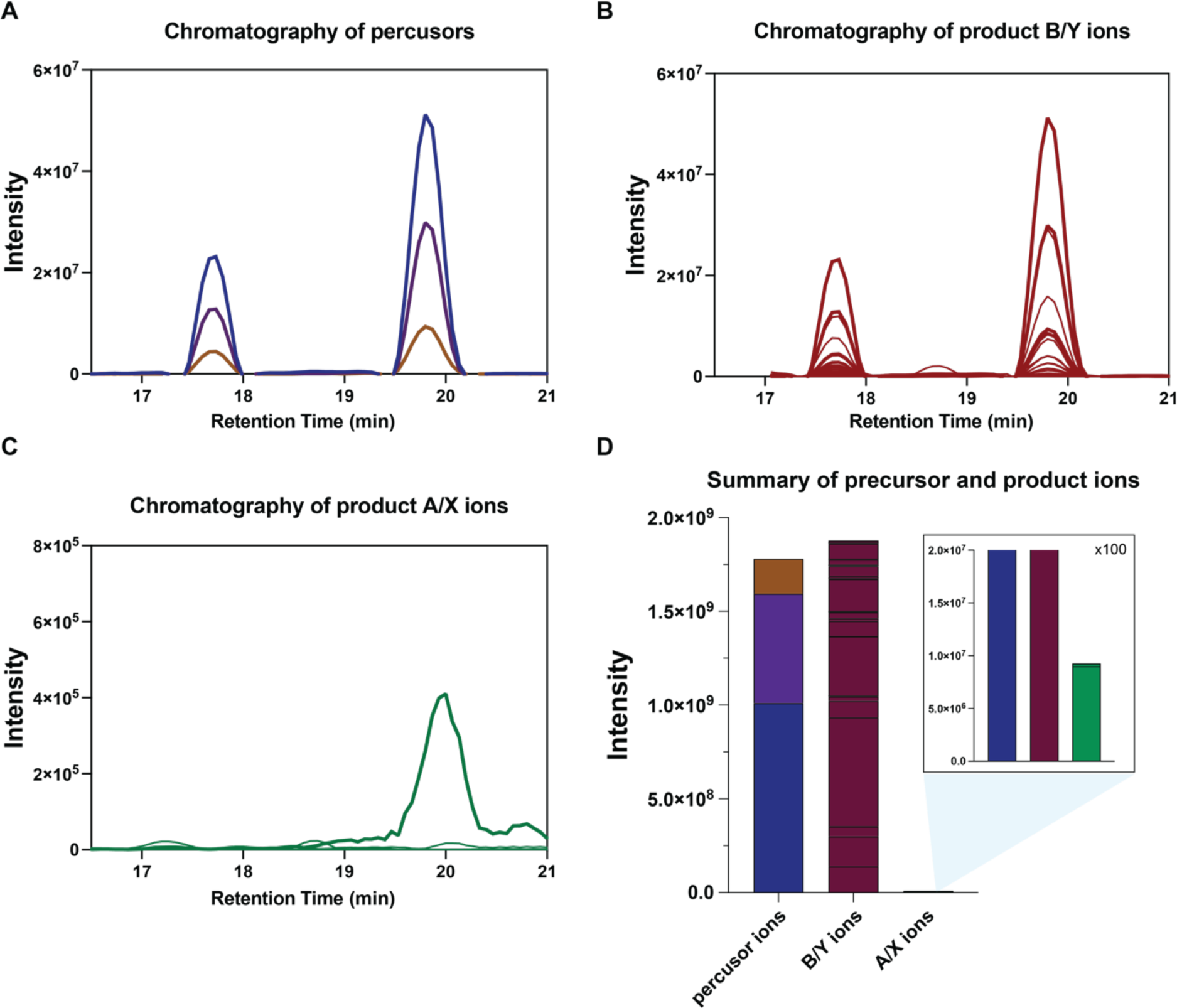
The chromatograms of example Hex(5)HexNAc(2) glycan. **A.** The chromatogram of precursor ions, including [M+] (blue), [M+1] (purple), and [M+2] (brown). **B.** The chromatogram of generated B, C, Y, and Z product ions from glycosidic bond cleavage. **C.** The chromatogram of generated A and X product ions from cross-ring dissociation. **D.** The summary of different ions from the example glycan showed more than 99.5% of product ions were generated from glycosidic bond cleavages at HCD NCE 20%.

**Figure S5.**
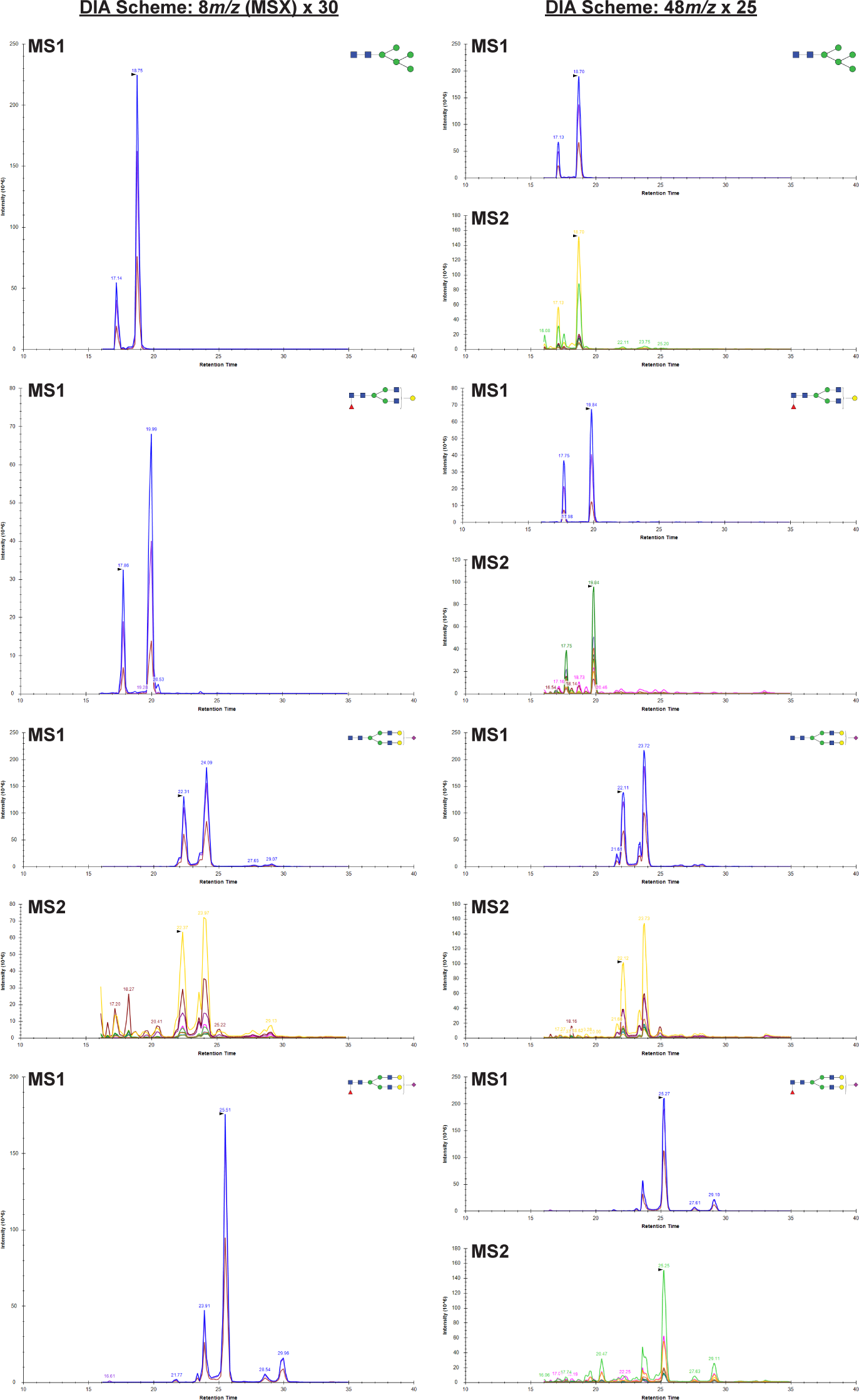
Evaluation of different DIA strategies and methods (part 1) Left, multiplexed (MSX) DIA strategy with 8 *m/z* isolation and 30 windows. Some of the glycan MS/MS information was missing due to the randomness of MSX selection. Right, fixed DIA strategy with 48 *m/z* isolation and 25 windows.

**Figure S6.**
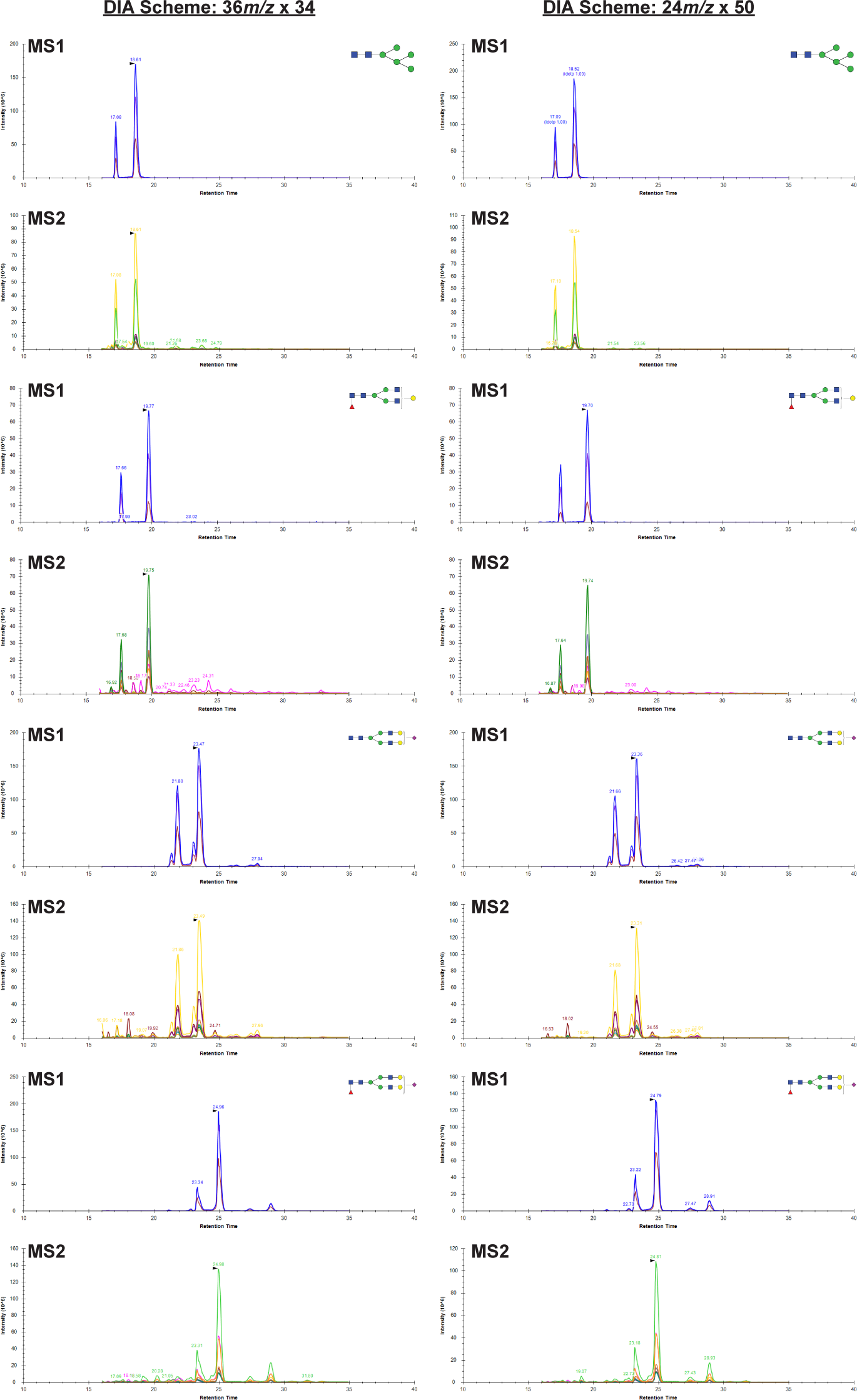
Evaluation of different DIA strategies and methods (part 2) Left, fixed DIA strategy with 36 *m/z* isolation and 34 windows. Right, fixed DIA strategy with 24 *m/z* isolation and 50 windows.

**Figure S7.**
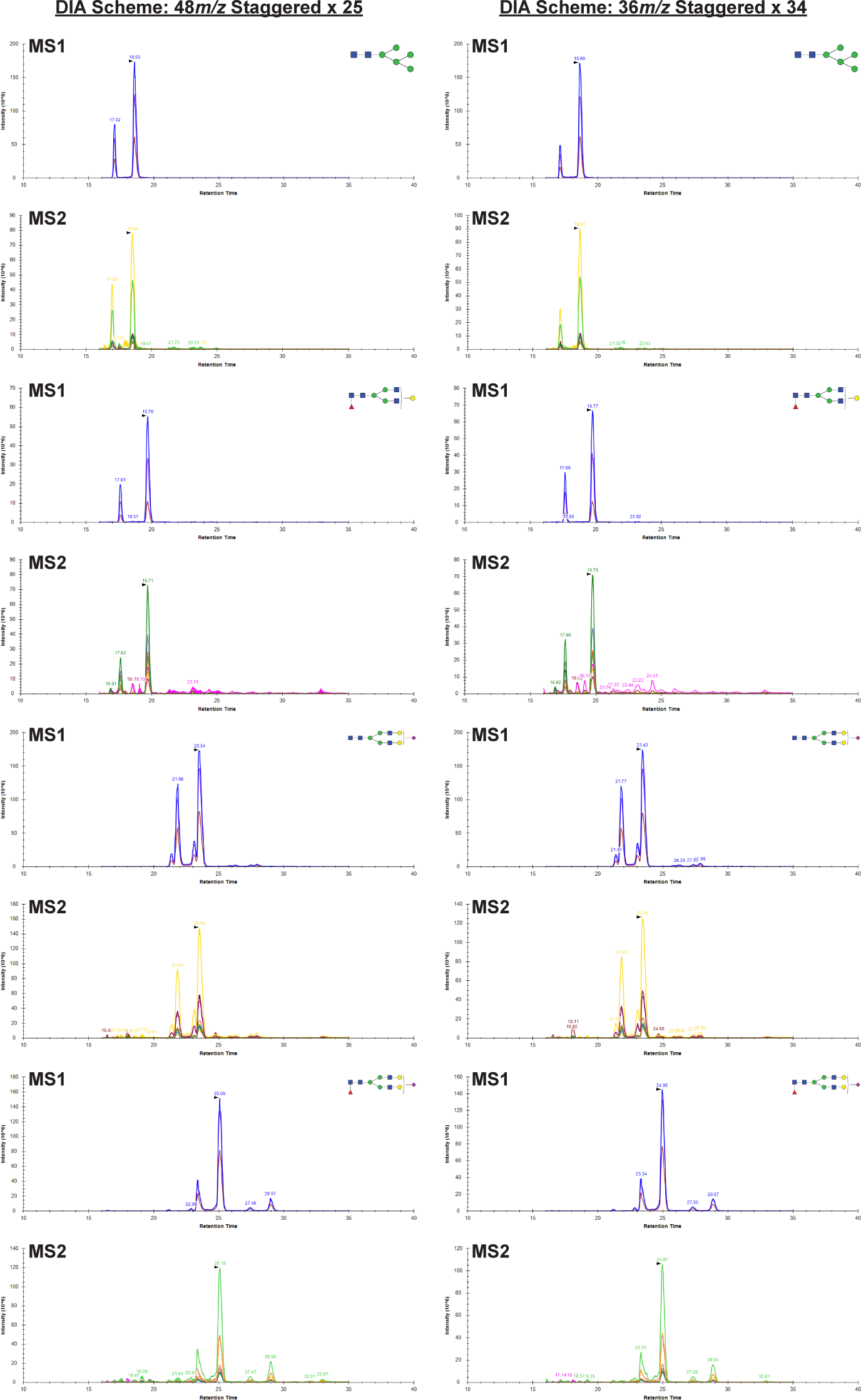
Evaluation of different DIA strategies and methods (part 3) Left, stagger DIA strategy with 48 *m/z* isolation and 25 windows. Right, stagger DIA strategy with 36 *m/z* isolation and 34 windows.

**Figure S8.**
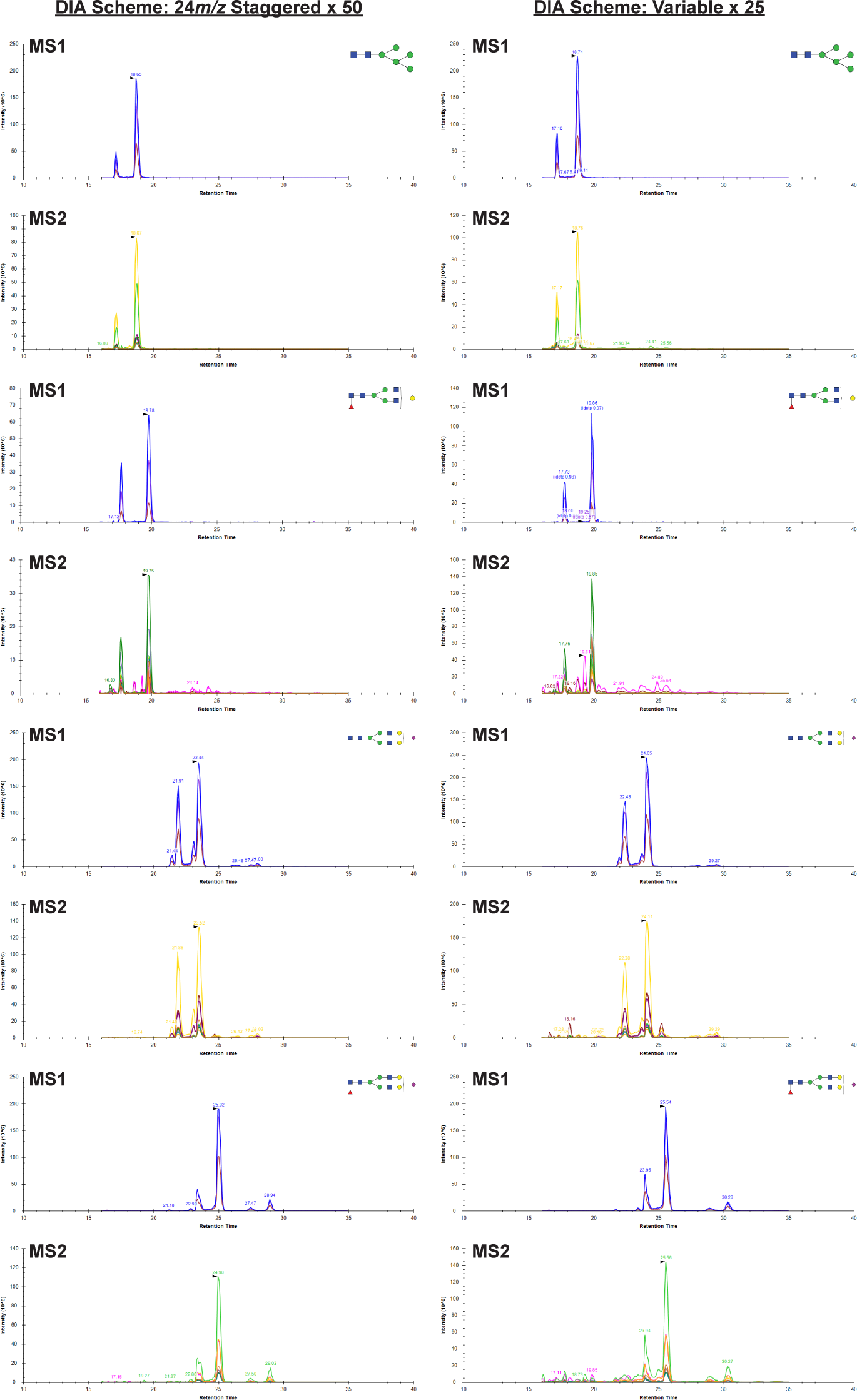
Evaluation of different DIA strategies and methods (part 4) Left, stagger DIA strategy with 24 *m/z* isolation and 50 windows. Right, variable DIA strategy with 25 windows, and the isolation window was calculated based on the ion distribution of analyzed glycans.

**Figure S9.**
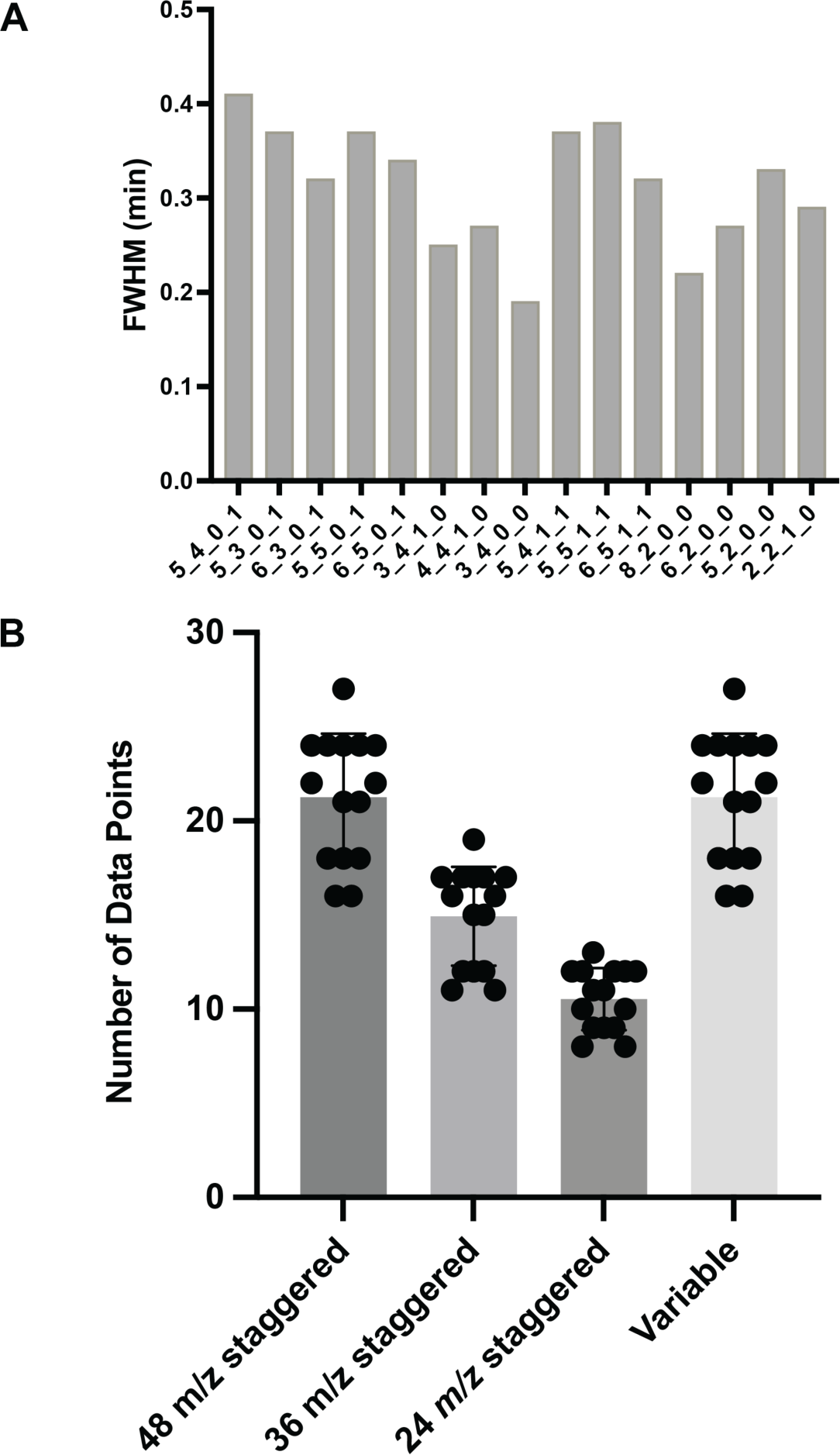
The full width at half maximum (FWHM) and data points across glycan peaks. **A.** In the PGC column, the average FWHM of glycans is more than 0.3 minutes, which provides enough length for DIA scans. **B.** Due to the retention of glycans in PGC materials, the GlycanDIA workflow enables more than 10 data points for different glycan peaks, which is optimal for accurate quantification.

**Figure S10.**
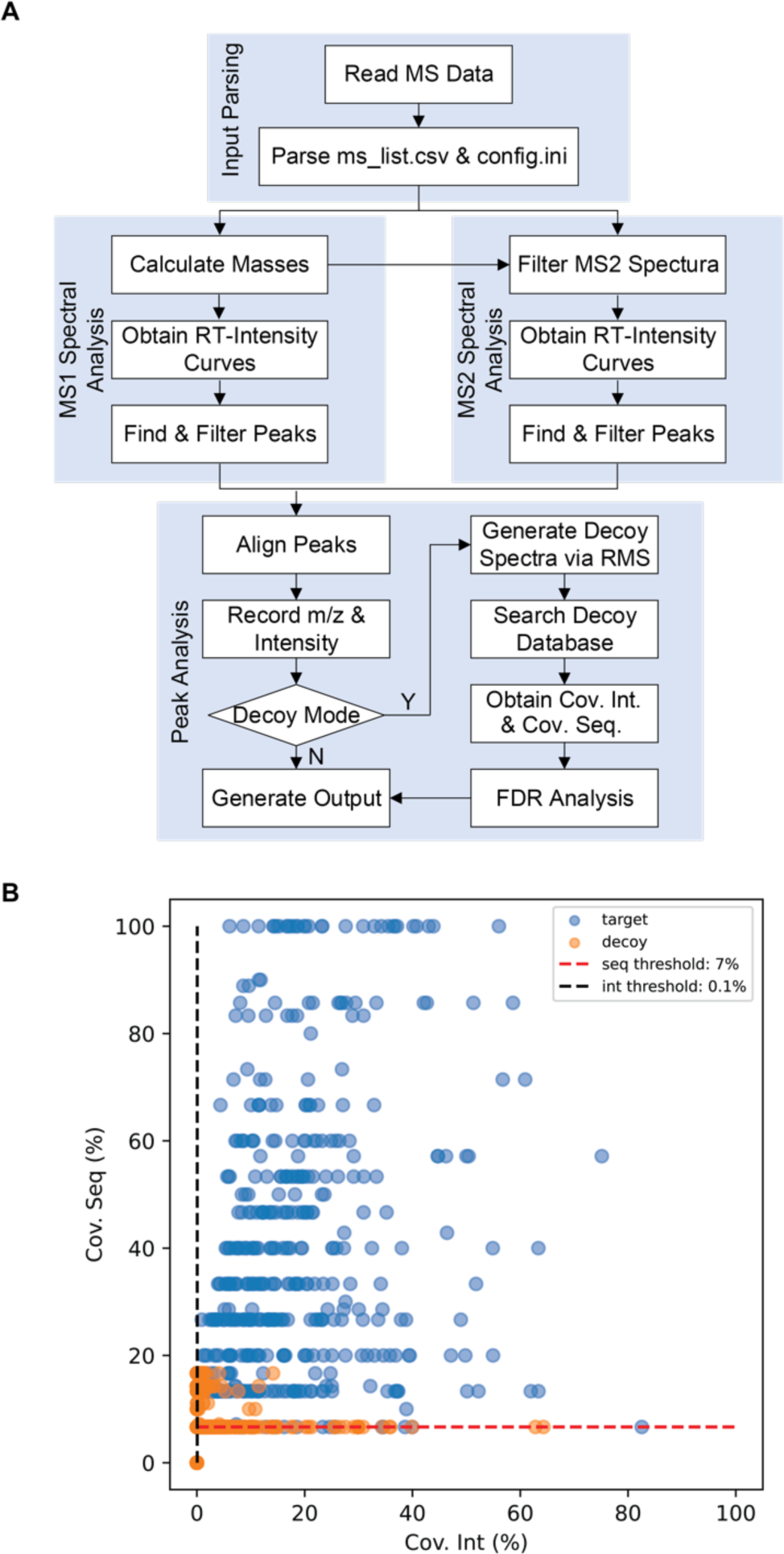
Data analysis using GlycanDIA Finder. **A.** GlycanDIA Finder is an open-source software tool built upon Python and MatchMS library to automatically search glycans and align peaks between the MS1 and MS2 data in a batch of MS files. The configuration settings, including input/output paths, polarity, adduct, charges, mass error, relative height and minimum height of peaks, the time window, and minimum matched count, control the behaviors of the software. The software also incorporated FDR calculation to remove the false identified glycans. **B.** Score distribution of target and decoy searches. Target search (blue) distributed in most parts of the density map, indicating that there are both high-quality and low-quality spectra. At the same time, spectra of decoy search(orange) distributed only in the low-left parts of the density map, indicating the distribution of false positives. The threshold is with 2% FDR. More details about the software can be found in **Supplementary Information 2.**

**Figure S11.**
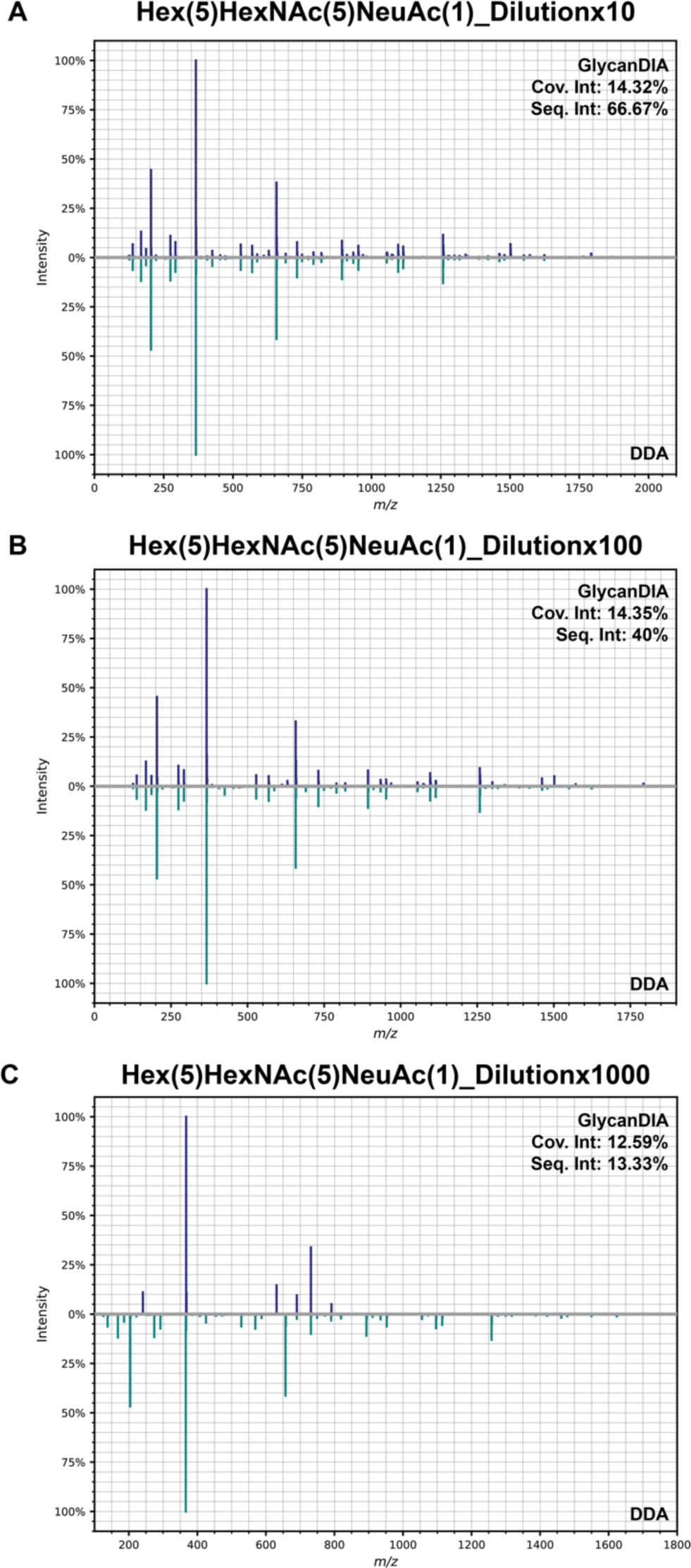
Head-to-tail comparison of sialylated glycan, Hex(5)HexNAc(5)Fuc(1) aligned spectra using GlycanDIA method against conventional DDA method. The aligned spectra showed a perfect match to the reference spectra up to 100-fold dilution, while user defined peaks were detected by GlycanDIA even after a 1000-fold dilution with a Cov. Int. 12.6% and Cov. Seq. 13.33%. The robustness of the alignment algorithm also indicates a potential library-free DIA analysis method for future development.

**Figure S12.**
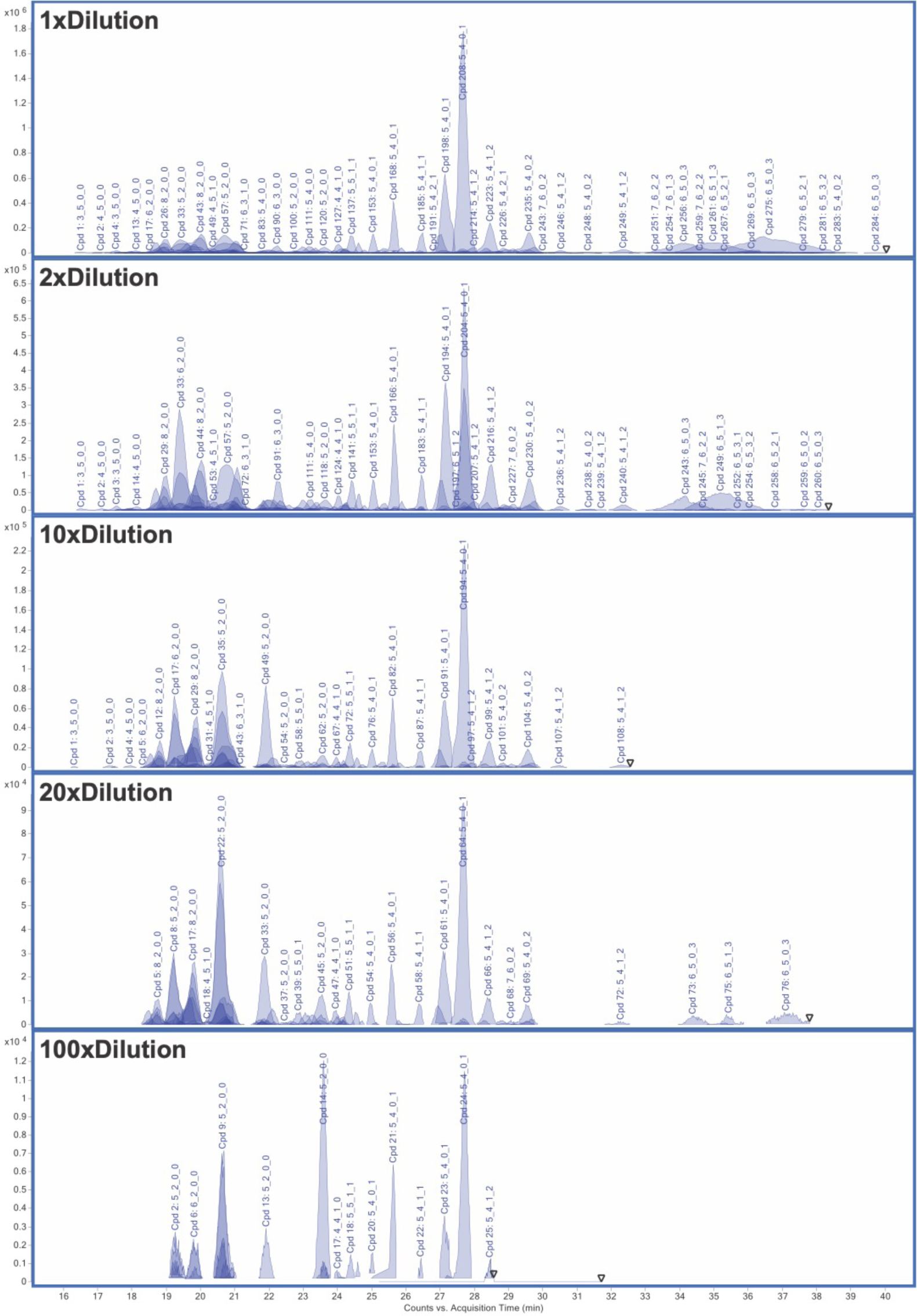
Characterization of glycans with different dilution factors using the nanoLC-ToF system. Each peak in the chromatogram represents a unique glycan compound (Cpd). The glycan peak can barely be quantified after a 100-time dilution of the glycan standard.

**Figure S13.**
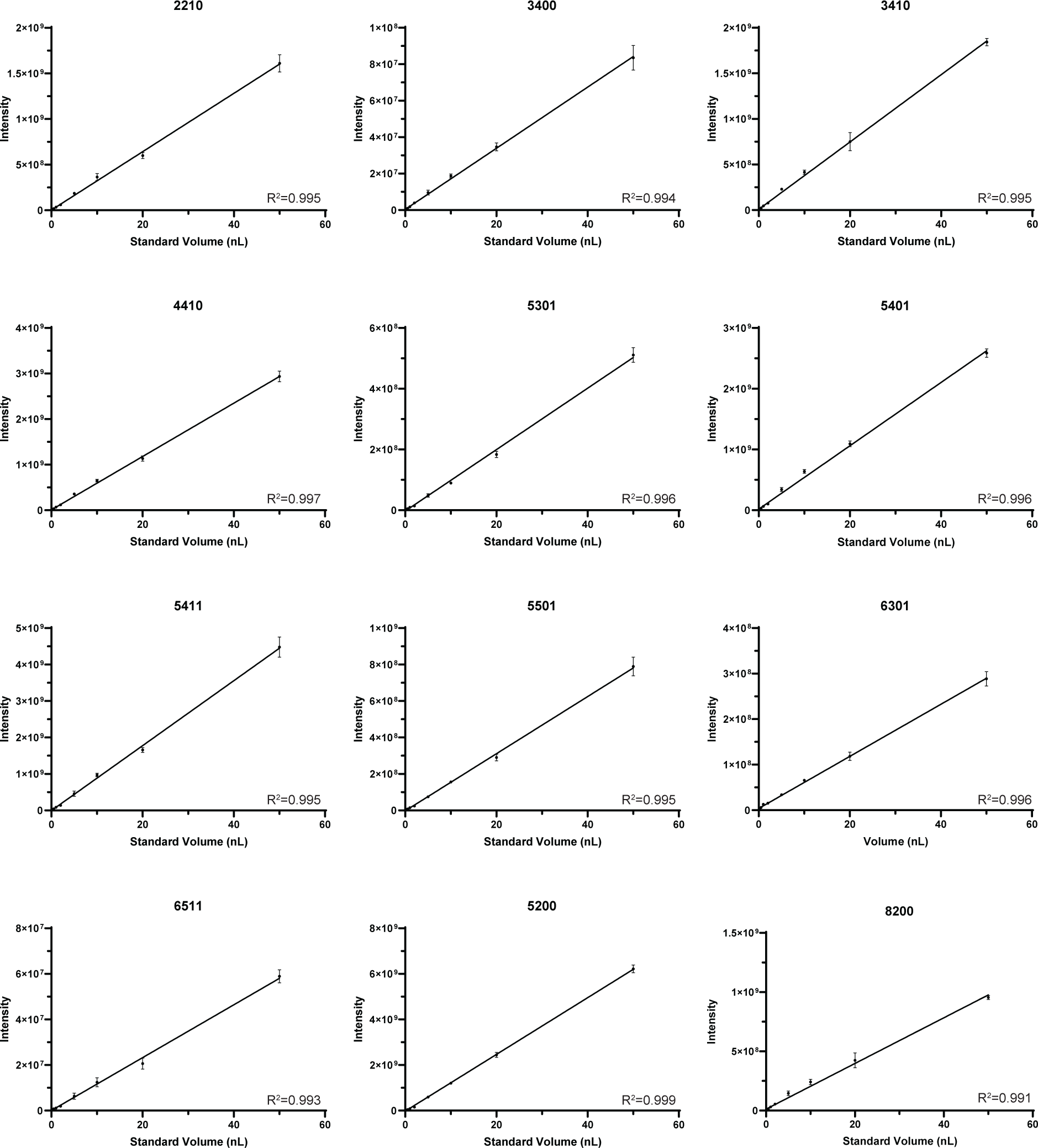
Calibration curves of different glycans using DIA-based nanoLC-orbitrap system. Unfilled points were not included in linear curves. All curves yielded superior *R*^2^ values (>0.99, and decent linearities were obtained after a 1000-time dilution for most glycans. These results demonstrated the sensitivity of the GlycanDIA workflow. Annotation is as follows: Hex_HexNAc_Fuc_Neu5Ac. demonstrated the sensitivity of the GlycanDIA workflow.

**Figure S14.**
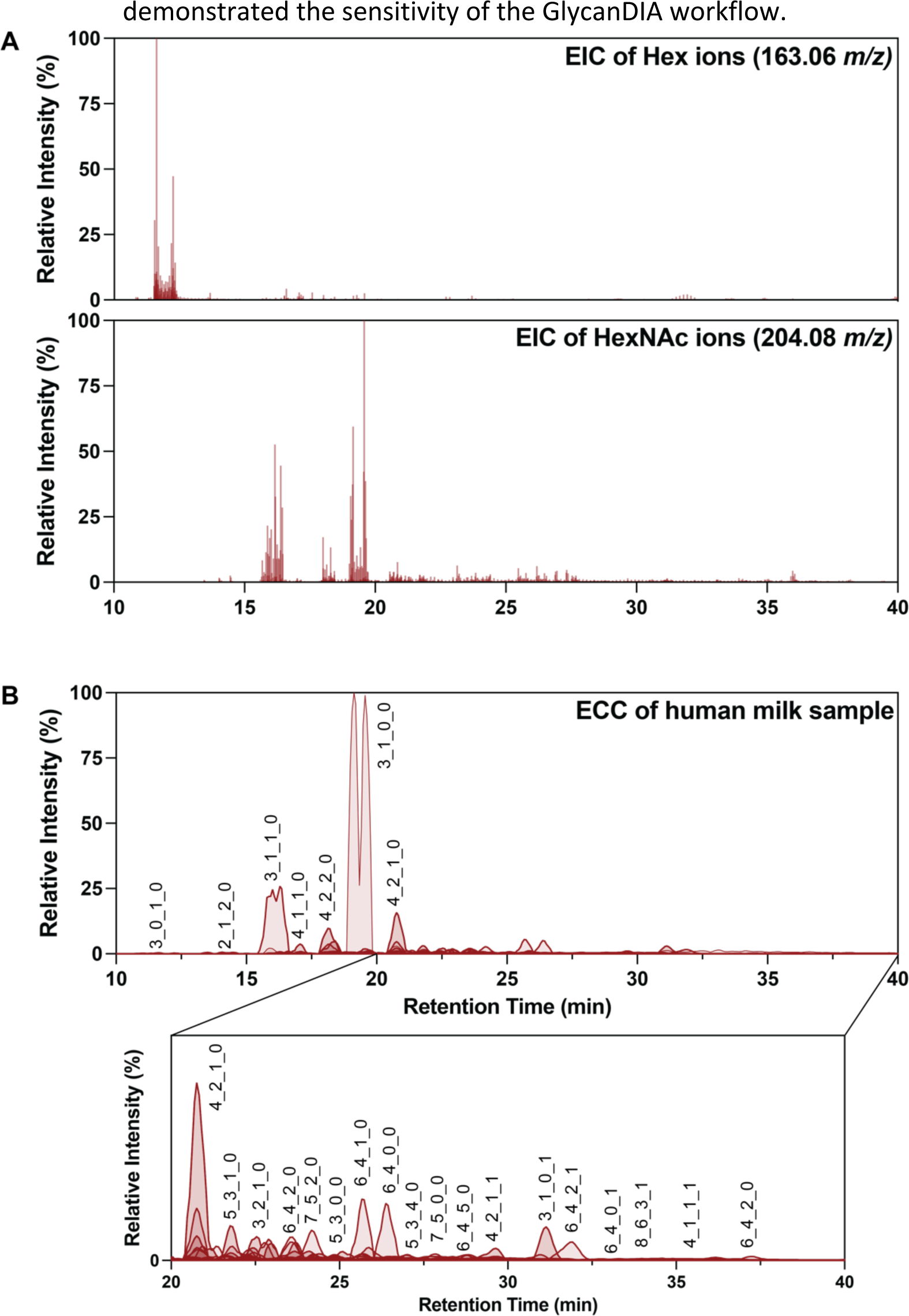
The profile of HMO from human milk sample using GlycanDIA. **A.** Extracted ion chromatogram (EIC) of 163.06 *m/z* and 204.08 *m/z* shows the signal of hexose- and N-acetylhexosamine-containing HMOs from the sample. **B.** Extracted compound chromatogram (ECC) shows the example elution profile of HMO from human milk. Annotation is as follows: Hex_HexNAc_Fuc_Neu5Ac.

**Figure S15.**
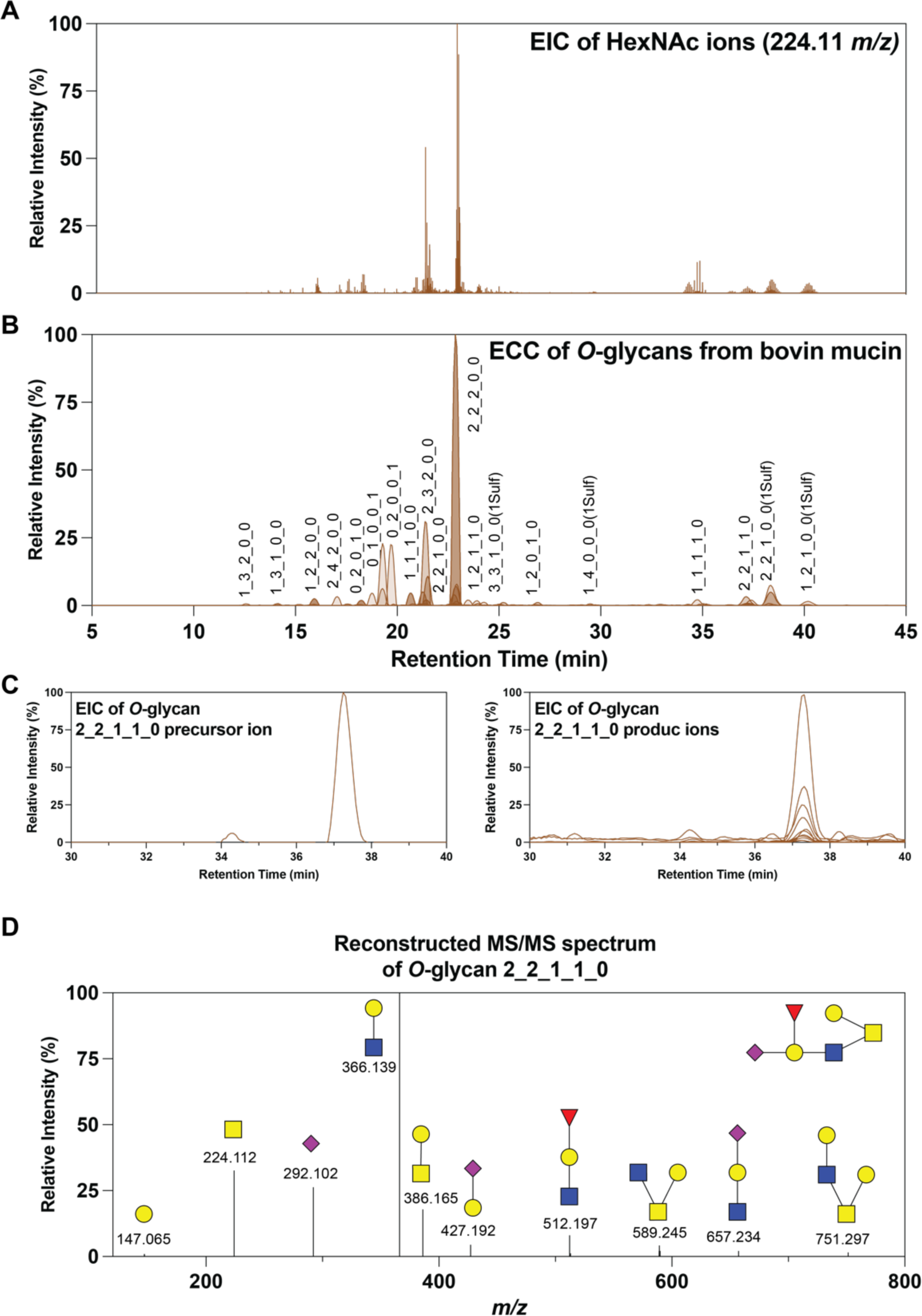
The profile of *O*-glycans from bovine mucin using GlycanDIA. **A.** EIC of 224.11 *m/z* shows the signal of reduced GalNAc. **B.** ECC shows the example elution profile of *O*-glycans from bovine mucin. Annotation is as follows: Hex_HexNAc_Fuc_Neu5Ac_Neu5Gc. **C.** The EIC of example *O*-glycan Hex(2)HexNAc(2)Fuc(1)Neu5Ac(1) precursor (left) and product (right) ions. **D.** The example annotation of reconstructed tandem mass spectrum of the example *O*-glycan. The fragment ions confirmed its structure, while the linkage specification is missed because HCD cannot reveal the general linkage information.

**Figure S16.**
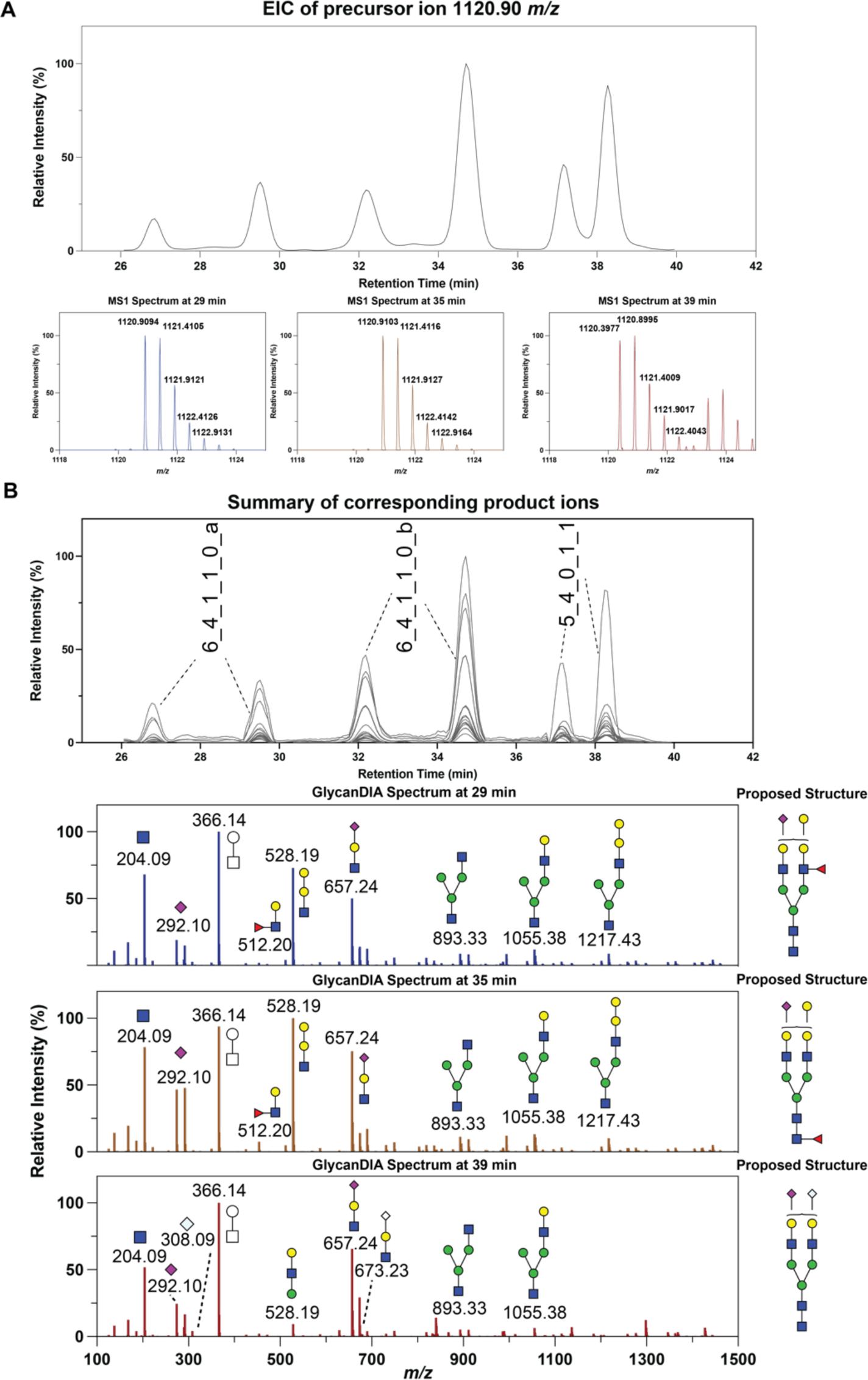
The example of identification of glycan composition isomers using GlycanDIA. **A.** Extracted ion chromatogram (EIC) of 1120.90 *m/z* at MS1 level, and six peaks corresponding to three different glycans and their anomers were noted. **B.** The extracted MS2 chromatogram of identified glycans and the representative glycan MS2 spectra at three different retention times, 29, 35, and 39min, with annotation and proposed structures.

**Figure S17.**
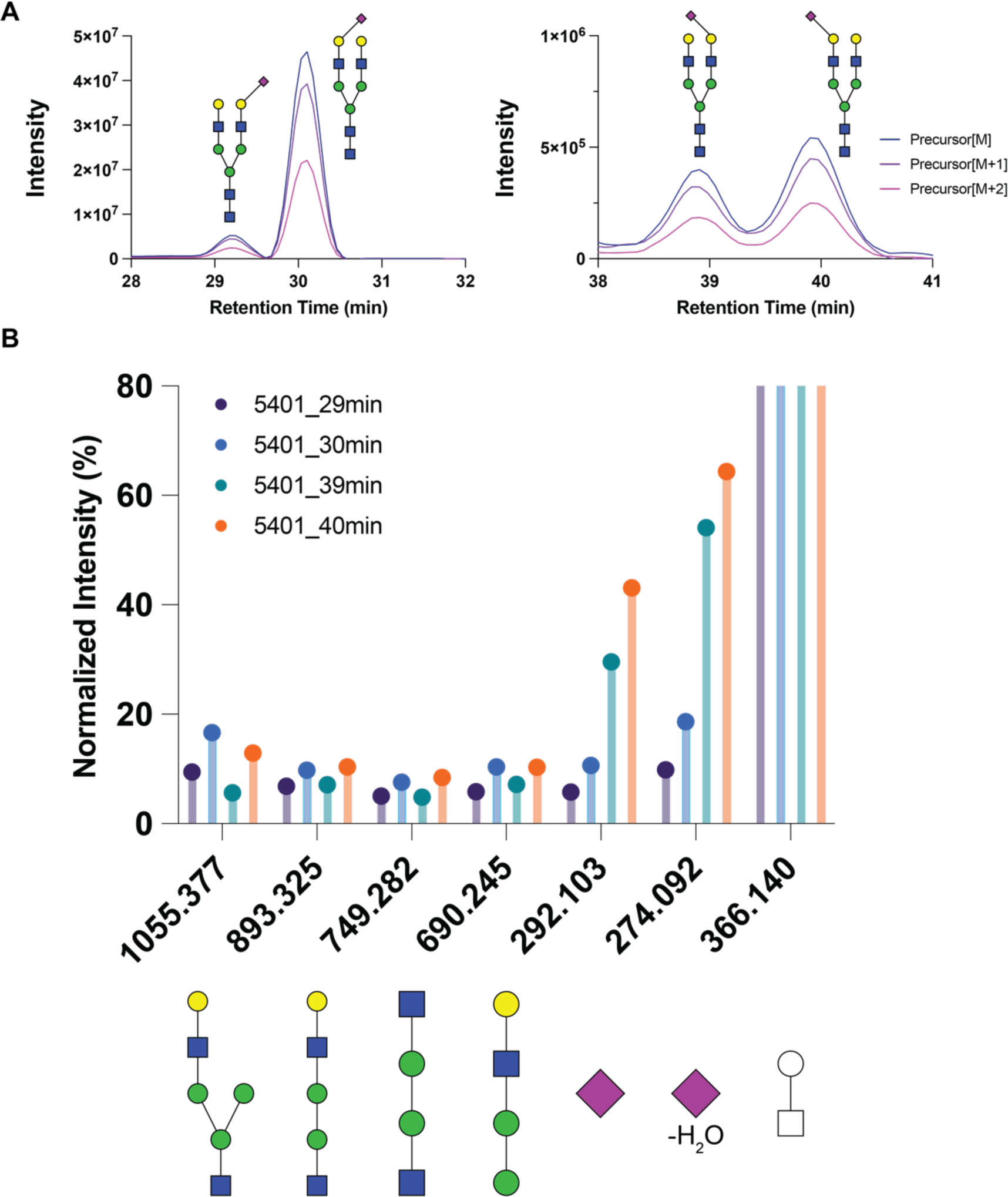
The example of identification of sialylated isomers using GlycanDIA. **A.** EIC of the glycan Man(3)Gal(2)GlcNAc(4)NeuAc(1) isomer precursor and its isotopic peaks at the MS1 level. **B.** The normalized intensity of different fragments from the example glycans. The product ion intensity was normalized to the fragment of 1Hexose+1HexNAc (366.14 *m/z*). Different product ion distributions reveal the potential capability to determine glycan linkage isomers using GlycanDIA. *Note: the specific glycan structure was based on the previous results reported by Palmisano *et al.* (*RSC Advances*, **2013**).

**Figure S18.**
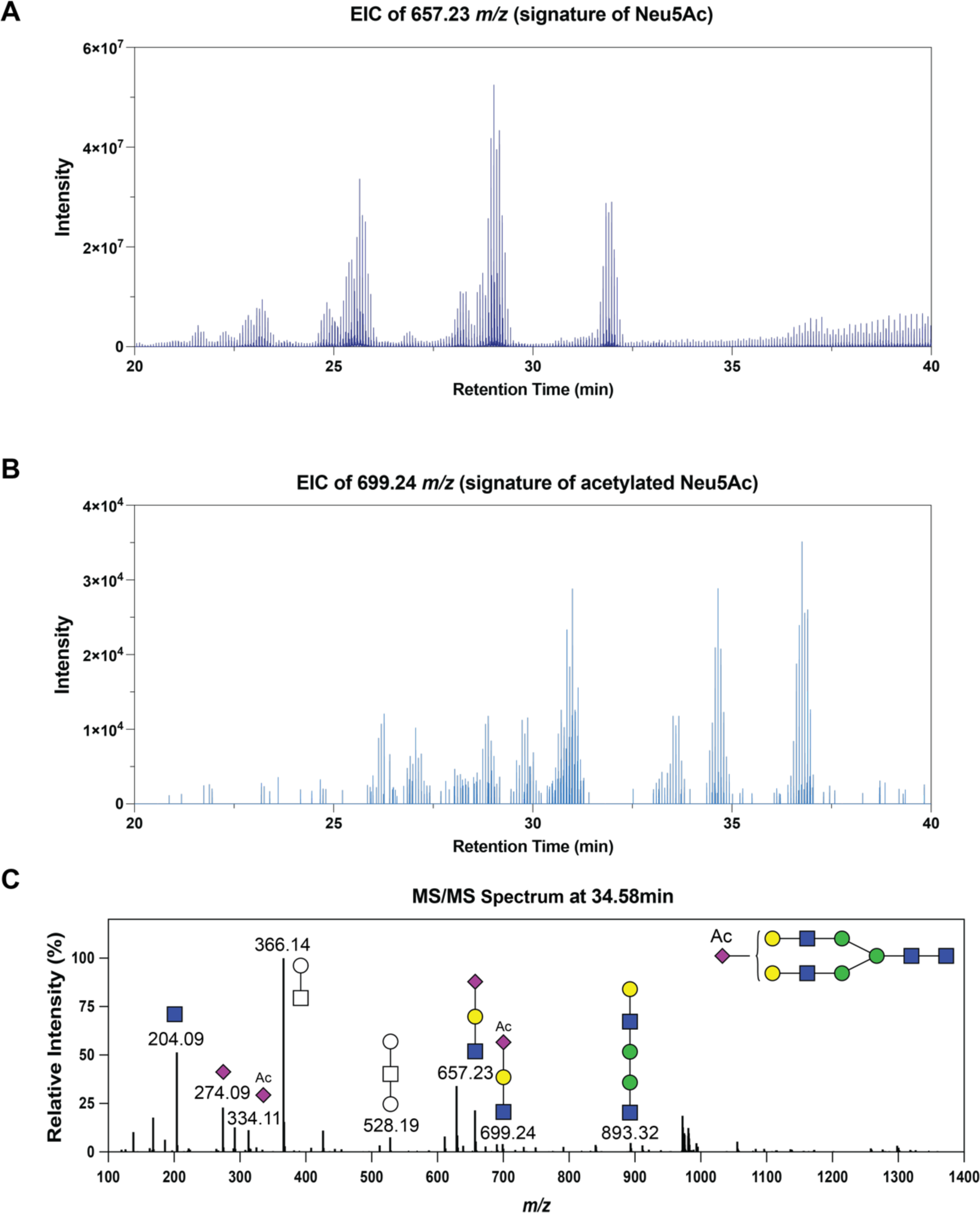
The identification of acetyl-modified glycans using GlycanDIA. **A.** EIC of 657.23 *m/z* representing the signature fragment from Neu5Ac-containing glycans. **B.** EIC of 699.24 *m/z* representing the signature fragment from acetylated Neu5Ac-containing glycans. **C.** The MS2 spectrum at 34.58min corresponding to an acetylated glycan, Hex(5)HexNAc(4)SiaAc(1), was shown as an example. The peak observed from the EIC demonstrates the low-abundant modified glycans can be identified using GlycanDIA.

**Figure S19.**
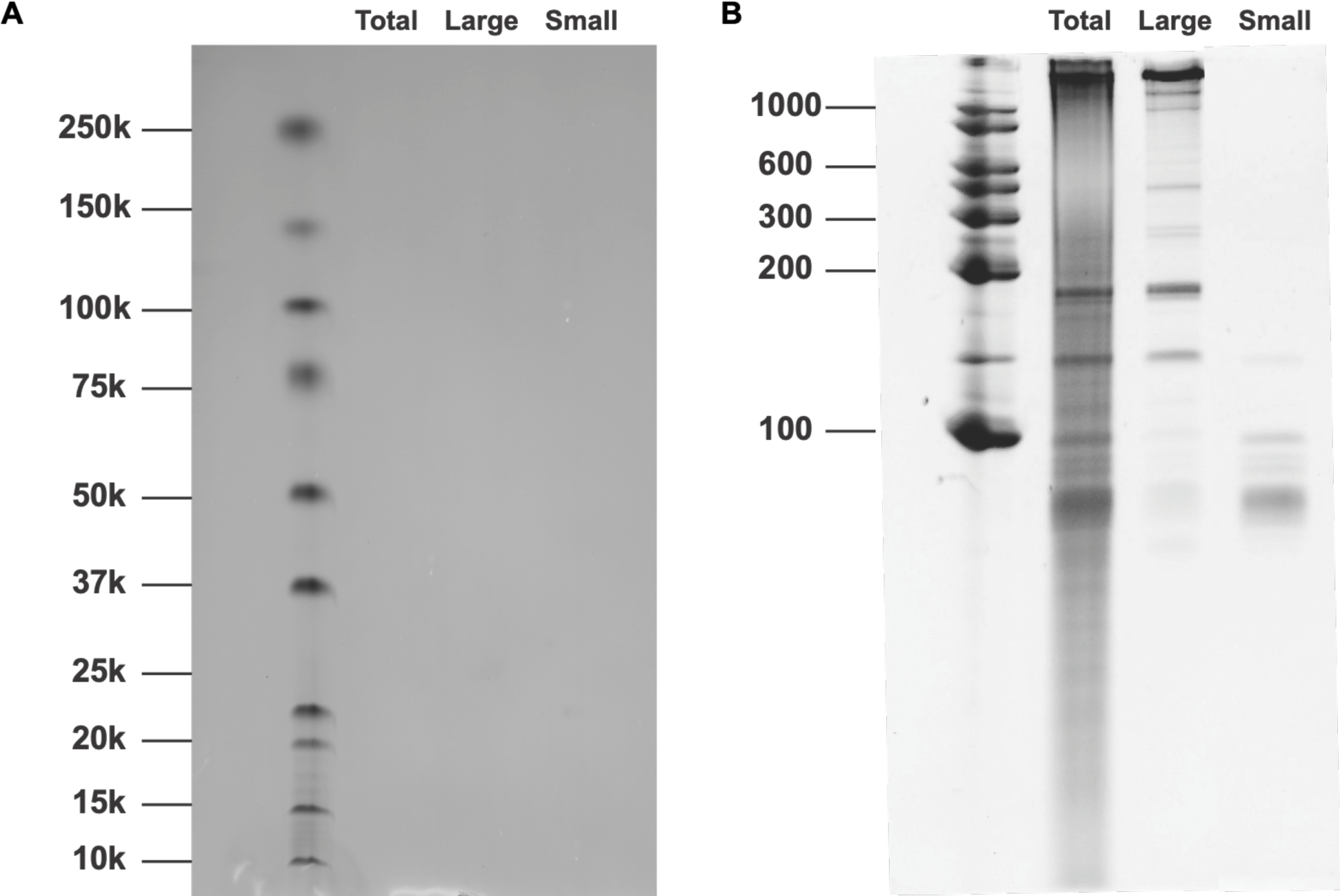
The protein and RNA gels of RNA samples. **A.** protein SDS-PAGE gel of the extracts. No protein band was observed from the RNA samples confirming the purity of RNA extracts. **B.** RNA TBE-urea gel of the extracts, the bands suggested successfully extraction of RNAs and the separation of large and small RNAs.

**Figure S20.**
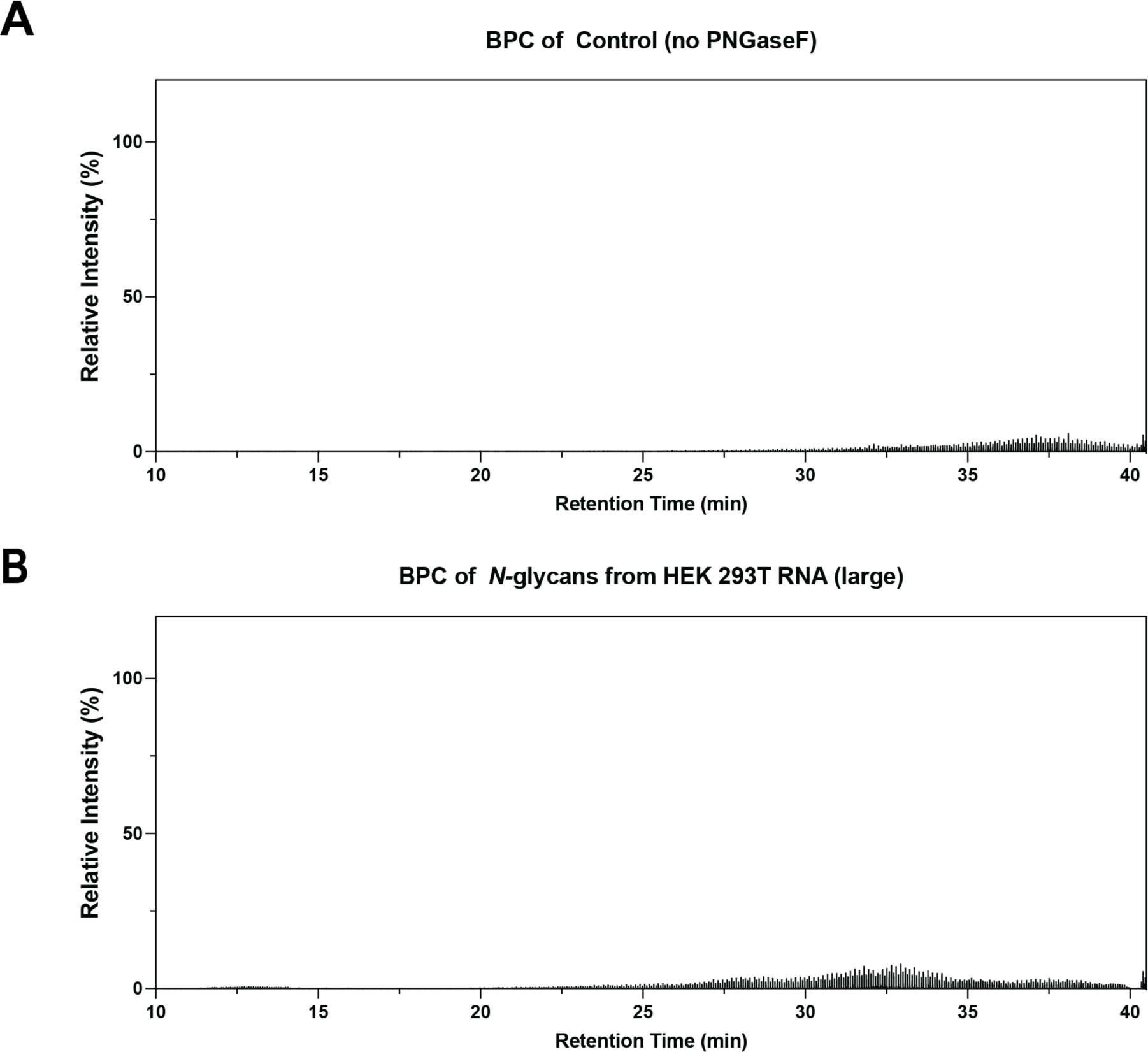
The chromatogram of *N*-glycans from HEK 293T cell line. **A.** The background signal from RNA extracts showed the limited of free glycans in the sample. **B.** Only background signals were noted from large RNA fractions, suggesting the glycosylation on RNA was barely in large RNAs.

**Figure S21.**
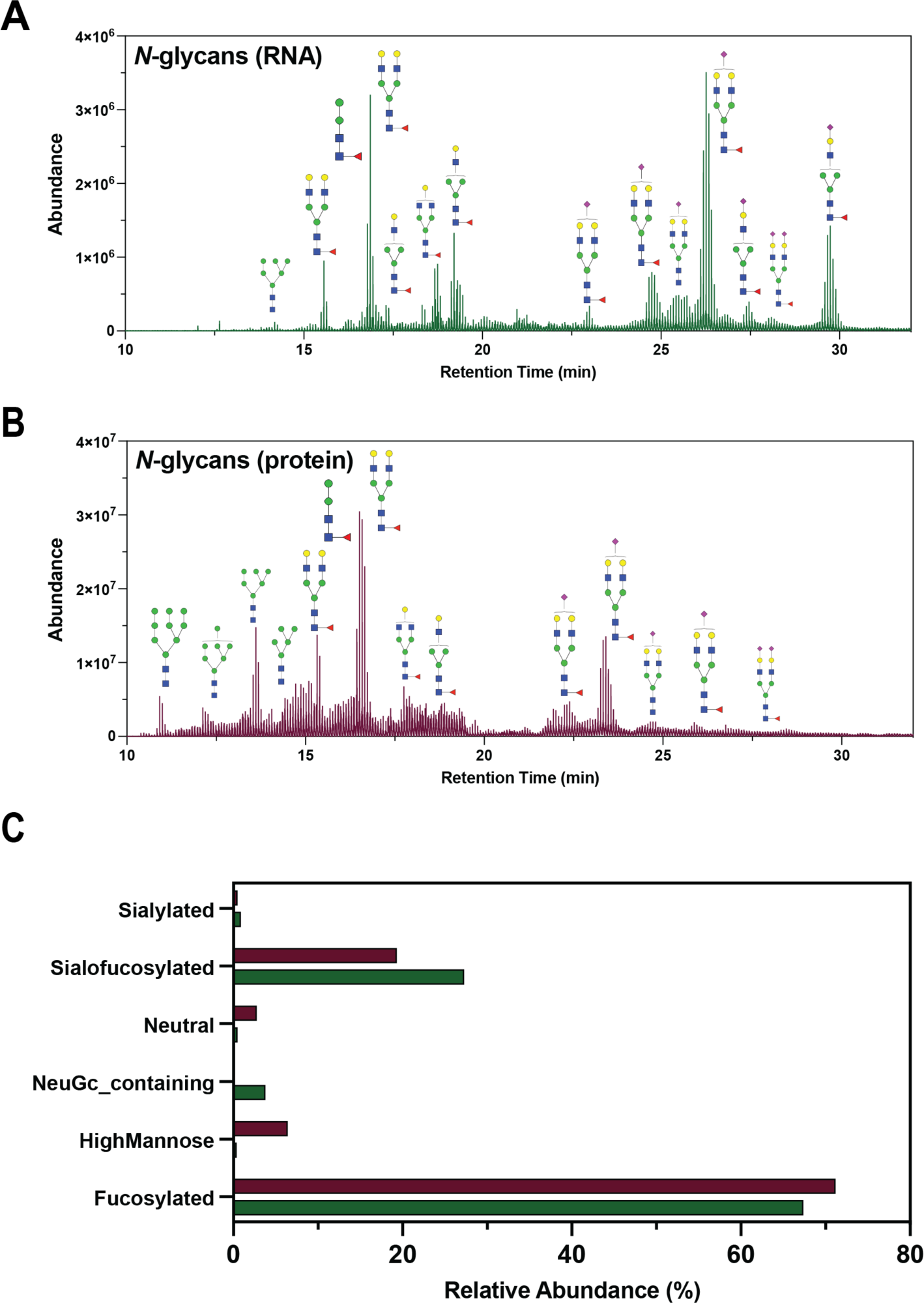
The chromatogram of *N*-glycans from HeLa cell line. **A.** Glycans from protein extracts showed higher abundance in both high mannose type and sialic acid-containing *N*-glycans. **B.** Glycans from small RNA fractions only showed higher abundance in the glycans with sialic acid. **C.** Only background signals were noted from large RNA fractions, suggesting the glycosylation on RNA was mainly in small RNAs. **C.** The relative abundance of RNA and protein *N*-glycans showed distinct glycan profiles.

**Figure S22.**
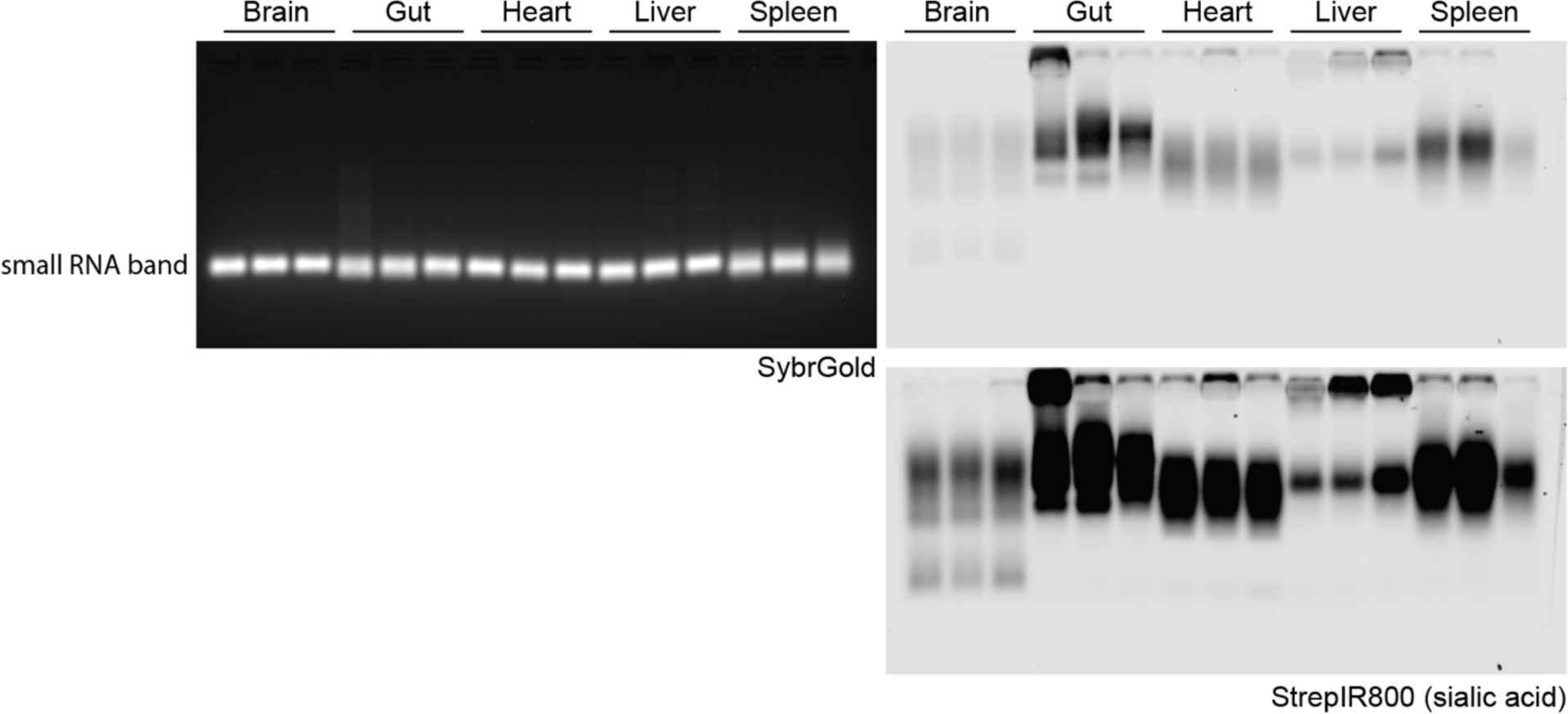
RNA blotting of small RNA from indicated mouse tissues labeled to detect the native sialic acids with rPAL. In gel detection of total RNA with SybrGold (left) and on membrane detection of biotin (Streptavidin-IR800, right) is shown. Two exposures of the biotin signal are displayed on the right. Each lane represents one biological replicate.

**Figure S23.**
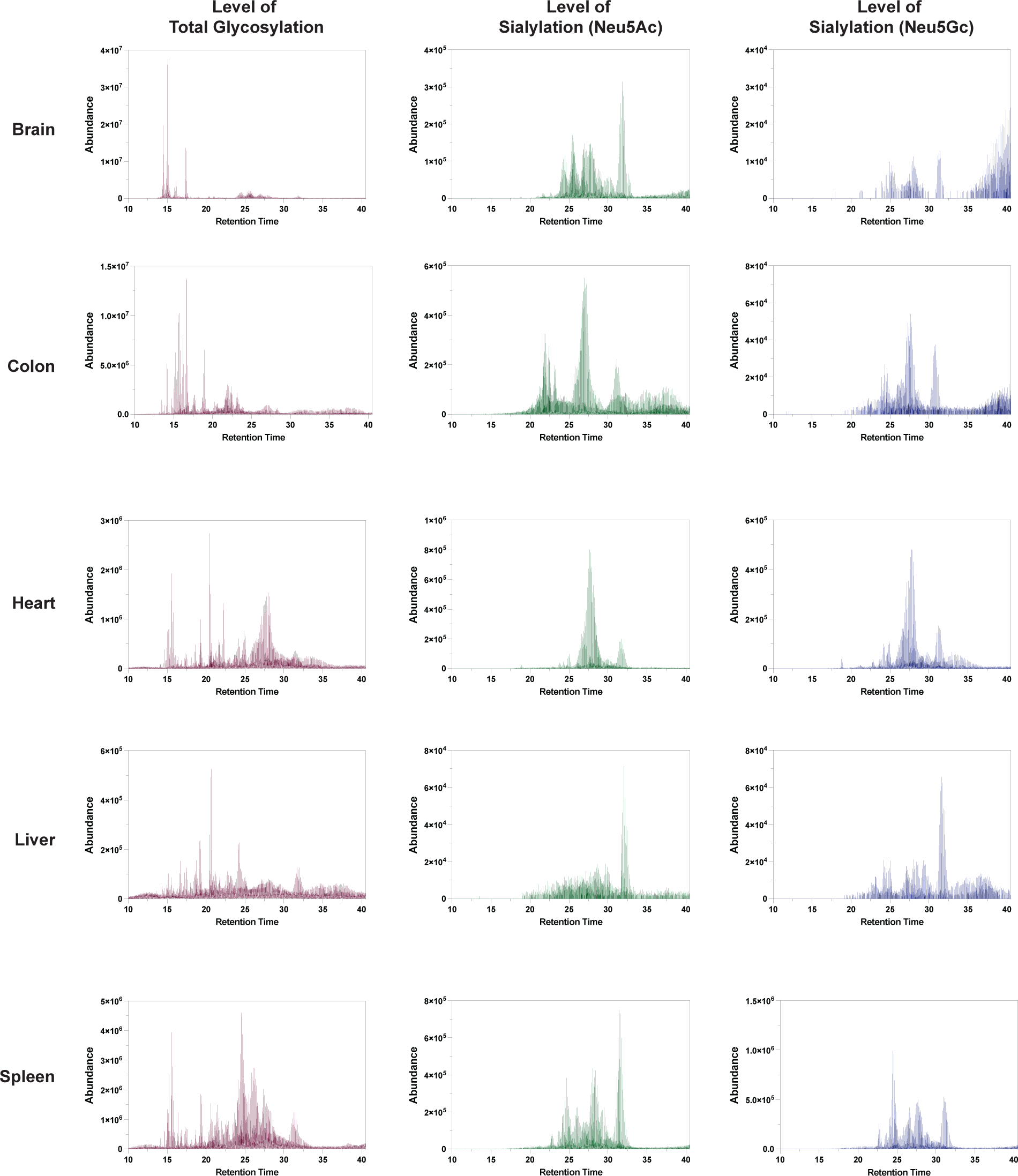
The level of Neu5Ac and Neu5Gc were expressed differently in mouse tissues. The glycan fragments of total glycosylation (red, 292.10 *m/z*), Neu5Ac-conatining glycans (green, 204.08 *m/z*), and Neu5Gc-contaning glycans (blue, 308.09 *m/z*).

## References

(1) Flynn, R. A.; Pedram, K.; Malaker, S. A.; Batista, P. J.; Smith, B. A. H.; Johnson, A. G.; George, B. M.; Majzoub, K.; Villalta, P. W.; Carette, J. E.; Bertozzi, C. R. Small RNAs are modified with N-glycans and displayed on the surface of living cells. Cell 2021, 184 (12), 3109–3124.e3122. DOI: 10.1016/j.cell.2021.04.023.

(2) Ajit Varki, R. D. C., Jeffrey D. Esko, Pamela Stanley, Gerald W. Hart, Markus Aebi, Debra Mohnen, Taroh Kinoshita, Nicolle H. Packer, James H. Prestegard, Ronald L. Schnaar, and Peter H. Seeberger. Essentials of Glycobiology, 4th *edition*; Cold Spring Harbor Laboratory Press, 2022.

(3) Xie, Y.; Hemberger, H.; Till, N. A.; Chai, P.; Watkins, C. P.; Lebedenko, C. G.; Caldwell, R. M.; George, B. M.; Bertozzi, C. R.; Garcia, B. A.; Flynn, R. A. The modified RNA base acp3U is an attachment site for N-glycans in glycoRNA. bioRxiv 2023, 2023.2011.2006.565735. DOI: 10.1101/2023.11.06.565735.

(4) Reily, C.; Stewart, T. J.; Renfrow, M. B.; Novak, J. Glycosylation in health and disease. Nature Reviews Nephrology 2019, 15 (6), 346–366. DOI: 10.1038/s41581-019-0129-4.

(5) Battistel, M. D.; Azurmendi, H. F.; Yu, B.; Freedberg, D. I. NMR of glycans: Shedding new light on old problems. Progress in Nuclear Magnetic Resonance Spectroscopy 2014, 79, 48–68. DOI: 10.1016/j.pnmrs.2014.01.001.

(6) Hirabayashi, J. Lectin-based structural glycomics: Glycoproteomics and glycan profiling. Glycoconjugate Journal 2004, 21 (1), 35–40. DOI: 10.1023/B:GLYC.0000043745.18988.a1.

(7) Ruhaak, L. R.; Xu, G.; Li, Q.; Goonatilleke, E.; Lebrilla, C. B. Mass Spectrometry Approaches to Glycomic and Glycoproteomic Analyses. Chemical Reviews 2018, 118 (17), 7886–7930. DOI: 10.1021/acs.chemrev.7b00732.

(8) Li, Q.; Xie, Y.; Wong, M.; Lebrilla, C. B. Characterization of Cell Glycocalyx with Mass Spectrometry Methods. Cells 2019, 8 (8), 882.

(9) Delafield, D. G.; Li, L. Recent Advances in Analytical Approaches for Glycan and Glycopeptide Quantitation. Molecular & Cellular Proteomics 2021, 20. DOI: 10.1074/mcp.R120.002095.

(10) Kasper, D. M.; Hintzen, J.; Wu, Y.; Ghersi, J. J.; Mandl, H. K.; Salinas, K. E.; Armero, W.; He, Z.; Sheng, Y.; Xie, Y.;, et al. The N-glycome regulates the endothelial-to-hematopoietic transition. Science 2020, 370 (6521), 1186–1191. DOI: 10.1126/science.aaz2121.

(11) Ruhaak, L. R.; Zauner, G.; Huhn, C.; Bruggink, C.; Deelder, A. M.; Wuhrer, M. Glycan labeling strategies and their use in identification and quantification. Analytical and Bioanalytical Chemistry 2010, 397 (8), 3457–3481. DOI: 10.1007/s00216-010-3532-z.

(12) Zhou, S.; Hu, Y.; DeSantos-Garcia, J. L.; Mechref, Y. Quantitation of Permethylated N-Glycans through Multiple-Reaction Monitoring (MRM) LC-MS/MS. Journal of the American Society for Mass Spectrometry 2015, 26 (4), 596–603. DOI: 10.1007/s13361-014-1054-1.

(13) Venable, J. D.; Dong, M.-Q.; Wohlschlegel, J.; Dillin, A.; Yates, J. R. Automated approach for quantitative analysis of complex peptide mixtures from tandem mass spectra. Nature Methods 2004, 1 (1), 39–45. DOI: 10.1038/nmeth705.

(14) Doerr, A. DIA mass spectrometry. Nature Methods 2015, 12 (1), 35–35. DOI: 10.1038/nmeth.3234.

(15) Raetz, M.; Bonner, R.; Hopfgartner, G. SWATH-MS for metabolomics and lipidomics: critical aspects of qualitative and quantitative analysis. Metabolomics 2020, 16 (6), 71. DOI: 10.1007/s11306-020-01692-0.

(16) Xie, Y.; De Luna Vitorino, F. N.; Chen, Y.; Lempiäinen, J. K.; Zhao, C.; Steinbock, R. T.; Lin, Z.; Liu, X.; Zahn, E.; Garcia, A. L.;, et al. SWAMNA: a comprehensive platform for analysis of nucleic acid modifications. Chemical Communications 2023, 59 (83), 12499–12502, 10.1039/D3CC04402E. DOI: 10.1039/D3CC04402E.

(17) De Leoz, M. L. A.; Simón-Manso, Y.; Woods, R. J.; Stein, S. E. Cross-Ring Fragmentation Patterns in the Tandem Mass Spectra of Underivatized Sialylated Oligosaccharides and Their Special Suitability for Spectrum Library Searching. Journal of the American Society for Mass Spectrometry 2019, 30 (3), 426–438. DOI: 10.1007/s13361-018-2106-8.

(18) Wong, M.; Xu, G.; Barboza, M.; Maezawa, I.; Jin, L.-W.; Zivkovic, A.; Lebrilla, C. B. Metabolic flux analysis of the neural cell glycocalyx reveals differential utilization of monosaccharides. Glycobiology 2020, 30 (11), 859–871. DOI: 10.1093/glycob/cwaa038.

(19) Xie, Y.; Chen, S.; Alvarez, M. R.; Sheng, Y.; Li, Q.; Maverakis, E.; Lebrilla, C. B. Protein oxidation of fucose environments (POFE) reveals fucose–protein interactions. Chemical Science 2024, 10.1039/D3SC06432H. DOI: 10.1039/D3SC06432H.

(20) Florian Huber, S. V., Christiaan Meijer, Hanno Spreeuw, Efraín Manuel Villanueva Castilla, Cunliang Geng, Justin J. J. van der Hooft, Simon Rogers, Adam Belloum, Faruk Diblen, and Jurriaan H. Spaaks1. matchms - processing and similarity evaluation of mass spectrometry data. Journal of Open Source Software 2020, 5 (52), 2411.

(21) Chen, Z.; Wei, J.; Tang, Y.; Lin, C.; Costello, C. E.; Hong, P. GlycoDeNovo2: An Improved MS/MS-Based De Novo Glycan Topology Reconstruction Algorithm. Journal of the American Society for Mass Spectrometry 2022, 33 (3), 436–445. DOI: 10.1021/jasms.1c00288.

(22) Liu, M.-Q.; Treves, G.; Amicucci, M.; Guerrero, A.; Xu, G.; Gong, T.-Q.; Davis, J.; Park, D.; Galermo, A.; Wu, L.;, et al. GlycoNote with Iterative Decoy Searching and Open-Search Component Analysis for High-Throughput and Reliable Glycan Spectral Interpretation. Analytical Chemistry 2023, 95 (21), 8223–8231. DOI: 10.1021/acs.analchem.3c00083.

(23) Thomsson, K. A.; Bäckström, M.; Holmén Larsson, J. M.; Hansson, G. C.; Karlsson, H. Enhanced Detection of Sialylated and Sulfated Glycans with Negative Ion Mode Nanoliquid Chromatography/Mass Spectrometry at High pH. Analytical Chemistry 2010, 82 (4), 1470–1477. DOI: 10.1021/ac902602e.

(24) Xu, G.; Davis, J. C. C.; Goonatilleke, E.; Smilowitz, J. T.; German, J. B.; Lebrilla, C. B. Absolute Quantitation of Human Milk Oligosaccharides Reveals Phenotypic Variations during Lactation1. The Journal of Nutrition 2017, 147 (1), 117–124. DOI: 10.3945/jn.116.238279.

(25) Xu, G.; Goonatilleke, E.; Wongkham, S.; Lebrilla, C. B. Deep Structural Analysis and Quantitation of O-Linked Glycans on Cell Membrane Reveal High Abundances and Distinct Glycomic Profiles Associated with Cell Type and Stages of Differentiation. Analytical Chemistry 2020, 92 (5), 3758–3768. DOI: 10.1021/acs.analchem.9b05103.

(26) Pett, C.; Nasir, W.; Sihlbom, C.; Olsson, B.-M.; Caixeta, V.; Schorlemer, M.; Zahedi, R. P.; Larson, G.; Nilsson, J.; Westerlind, U. Effective Assignment of α2,3/α2,6-Sialic Acid Isomers by LC-MS/MS-Based Glycoproteomics. Angewandte Chemie International Edition 2018, 57 (30), 9320–9324. DOI: 10.1002/anie.201803540.

(27) Palmisano, G.; Larsen, M. R.; Packer, N. H.; Thaysen-Andersen, M. Structural analysis of glycoprotein sialylation – part II: LC-MS based detection. RSC Advances 2013, 3 (45), 22706–22726, 10.1039/C3RA42969E. DOI: 10.1039/C3RA42969E.

(28) Muthana, S. M.; Campbell, C. T.; Gildersleeve, J. C. Modifications of Glycans: Biological Significance and Therapeutic Opportunities. ACS Chemical Biology 2012, 7 (1), 31–43. DOI: 10.1021/cb2004466.

(29) Haltiwanger, R. S. Fucose Is on the TRAIL of Colon Cancer. Gastroenterology 2009, 137 (1), 36–39. DOI: 10.1053/j.gastro.2009.05.010.

(30) Johannes, H.; Stefan, M.; Tiago, O.; Josef, M. P.; Friedrich, A.; Johannes, S. The Mouse N-Glycome Atlas – High-resolution N-glycan analysis of 23 mouse tissues. bioRxiv 2022, 2022.2009.2019.508483. DOI: 10.1101/2022.09.19.508483.

(31) Otaki, M.; Hirane, N.; Natsume-Kitatani, Y.; Nogami Itoh, M.; Shindo, M.; Kurebayashi, Y.; Nishimura, S.-I. Mouse tissue glycome atlas 2022 highlights inter-organ variation in major N-glycan profiles. Scientific Reports 2022, 12 (1), 17804. DOI: 10.1038/s41598-022-21758-4.

(32) Helena, H.; Peiyuan, C.; Charlotta, G. L.; Reese, M. C.; Benson, M. G.; Ryan, A. F. Rapid and sensitive detection of native glycoRNAs. bioRxiv 2023, 2023.2002.2026.530106. DOI: 10.1101/2023.02.26.530106.

(33) Stincone, P.; Pakkir Shah, A. K.; Schmid, R.; Graves, L. G.; Lambidis, S. P.; Torres, R. R.; Xia, S.-N.; Minda, V.; Aron, A. T.; Wang, M.;, et al. Evaluation of Data-Dependent MS/MS Acquisition Parameters for Non-Targeted Metabolomics and Molecular Networking of Environmental Samples: Focus on the Q Exactive Platform. Analytical Chemistry 2023, 95 (34), 12673–12682. DOI: 10.1021/acs.analchem.3c01202.

(34) Cho, B. G.; Gutierrez Reyes, C. D.; Goli, M.; Gautam, S.; Banazadeh, A.; Mechref, Y. Targeted N-Glycan Analysis with Parallel Reaction Monitoring Using a Quadrupole-Orbitrap Hybrid Mass Spectrometer. Analytical Chemistry 2022, 94 (44), 15215–15222. DOI: 10.1021/acs.analchem.2c01975.

(35) Wuhrer, M.; Koeleman, C. A. M.; Hokke, C. H.; Deelder, A. M. Mass spectrometry of proton adducts of fucosylated N-glycans: fucose transfer between antennae gives rise to misleading fragments. Rapid Communications in Mass Spectrometry 2006, 20 (11), 1747–1754. DOI: 10.1002/rcm.2509.

(36) Wuhrer, M.; Deelder, A. M.; van der Burgt, Y. E. M. Mass spectrometric glycan rearrangements. Mass Spectrometry Reviews 2011, 30 (4), 664–680. DOI: 10.1002/mas.20337.

(37) Wei, J.; Papanastasiou, D.; Kosmopoulou, M.; Smyrnakis, A.; Hong, P.; Tursumamat, N.; Klein, J. A.; Xia, C.; Tang, Y.; Zaia, J.;, et al. De novo glycan sequencing by electronic excitation dissociation MS2-guided MS3 analysis on an Omnitrap-Orbitrap hybrid instrument. Chemical Science 2023, 14 (24), 6695–6704, 10.1039/D3SC00870C. DOI: 10.1039/D3SC00870C.

(38) Ashwood, C.; Pratt, B.; MacLean, B. X.; Gundry, R. L.; Packer, N. H. Standardization of PGC-LC-MS-based glycomics for sample specific glycotyping. Analyst 2019, 144 (11), 3601–3612, 10.1039/C9AN00486F. DOI: 10.1039/C9AN00486F.

(39) Li, Q.; Xie, Y.; Wong, M.; Barboza, M.; Lebrilla, C. B. Comprehensive structural glycomic characterization of the glycocalyxes of cells and tissues. Nature Protocols 2020, 15 (8), 2668–2704. DOI: 10.1038/s41596-020-0350-4.

(40) Wu, S.; Tao, N.; German, J. B.; Grimm, R.; Lebrilla, C. B. Development of an Annotated Library of Neutral Human Milk Oligosaccharides. Journal of Proteome Research 2010, 9 (8), 4138–4151. DOI: 10.1021/pr100362f.

(41) Kessner, D.; Chambers, M.; Burke, R.; Agus, D.; Mallick, P. ProteoWizard: open source software for rapid proteomics tools development. Bioinformatics 2008, 24 (21), 2534–2536. DOI: 10.1093/bioinformatics/btn323.

(42) Ceroni, A.; Maass, K.; Geyer, H.; Geyer, R.; Dell, A.; Haslam, S. M. GlycoWorkbench: A Tool for the Computer-Assisted Annotation of Mass Spectra of Glycans. Journal of Proteome Research 2008, 7 (4), 1650–1659. DOI: 10.1021/pr7008252.

(43) Adams, K. J.; Pratt, B.; Bose, N.; Dubois, L. G.; St. John-Williams, L.; Perrott, K. M.; Ky, K.; Kapahi, P.; Sharma, V.; MacCoss, M. J.;, et al. Skyline for Small Molecules: A Unifying Software Package for Quantitative Metabolomics. Journal of Proteome Research 2020, 19 (4), 1447–1458. DOI: 10.1021/acs.jproteome.9b00640.

